# Unsupervised learning of mapping between brain lesions and behavior

**DOI:** 10.1101/2023.12.22.573110

**Authors:** Iman A. Wahle, Joseph Griffis, Ralph Adolphs, Jordan Grafman, Daniel Tranel, Aaron Boes, Frederick Eberhardt

## Abstract

Human lesion studies offer one of the most direct routes to investigating the relations between brain regions and behavioral outcomes in circumstances where experimental interventions are highly restricted. However, these studies face a major challenge in identifying the right level of granularity at which brain regions and behavioral outcomes should be analyzed to identify the relation between lesions to specific brain regions and specific behavioral outcomes. Here we showcase a novel data-driven approach, Causal Feature Learning (CFL), that learns the appropriate level of analysis and the relation between lesion and cognitive impairment at the same time. The method avoids specifying brain regions and specific outcome measures a priori, allowing for the discovery of new cross-cutting lesion-behavior maps. We show that CFL robustly recovers lesion behavior maps in a simulated dataset where Canonical Correlation Analysis fails to provide interpretable results. We then show that CFL recovers known lesion-behavior maps for language deficits and visuospatial processing using a large dataset of lesion subjects, and we illustrate how CFL can be used to identify new groupings of outcomes when mapping lesions to depression symptoms.

## Introduction

Ever since the seminal work of Broca, Wernicke, and others [1], demonstrably large and often remarkably specific cognitive impairments have been documented following macroscopic but focal brain lesions. Such case studies launched much of modern cognitive neuroscience [2, 3] and were the first to suggest that specific neural structures might be most essential for memory [4], aspects of face perception [5, 6], conscious vision [7], and complex decision-making [8]. Since the advent of CT and structural MRI, larger samples of subjects and more accurate methods of lesion localization have provided considerably more detailed, and more statistically reliable associations between lesion locations and specific cognitive impairments in domains ranging from facial emotion recognition [9] to aspects of language [10], object recognition [11], intelligence [12], and many other domains.

Human lesion studies offer a route to investigating the causal relations between brain regions and behavioral outcomes, though do so in the absence of controlled experimental interventions that are common in animal studies. Their interventional nature, even if accidents of nature rather than experimentally controlled, makes the results less prone to suffering from confounding, a concern often raised about similar findings based on large-scale functional neuroimaging studies in healthy subjects [13, 14]. However, the inference to a causal conclusion in lesion studies still faces the double challenge of establishing the cause-effect relation while simultaneously determining the causal *relata*, i.e. the quantities that stand in the causal relation: at what level of granularity should brain regions and behavioral outcomes be analyzed to identify the effects of specific brain regions on specific behavioral outcomes [15, 16]? Identifying such brain regions is crucial to the development of novel therapeutic treatments, but over-aggregation or mis-aggregation of voxel-level measurements can obscure the true causal relations present in the data. Simply put: Given a brain lesion and change in task performance, how can the lesion-to-behavior association best be characterized to support a generalizable causal claim?

Current approaches to lesion-behavior mapping typically involve associating single or multiple voxels with a single behavioral variable. Mass-univariate methods test each voxel independently for its association with a behavioral variable [17], while multivariate-univariate methods such as support vector regression [18] or sparse canonical correlation analysis (SCCAN) [19] instead model the relationship between the entire distribution of lesion location and a behavioral variable. While canonical correlation analysis (CCA), the model underlying SCCAN, can in principle be used with multi-dimensional behavioral variables, in practice this is rarely done in the context of lesion-behavior mapping due to challenges in interpretation (with a couple exceptions [20, 21]). Instead, multivariate behavioral measures are typically first reduced to a handful of summary variables using methods like factor analysis or principal component analysis, which are then related to multivariate lesion data one by one [22, 23, 24]. As a result, existing approaches typically consider behavioral measures independently of one another, and do so at a relatively coarse granularity by combining fine-grained measurements into single summary variables. Subsequently, lesion-behavior maps are often reduced to a binary association, labeling regions as either involved or not, when it is well-known that different brain regions may contribute to a behavior in distinct ways. These points strike us as an enormous missed opportunity: Key advances in cognition are likely to come from avoiding a priori restrictions on outcome measures and allowing lesion analyses to reveal previously unrecognized behavioral dissociations – behaviors tightly linked to specific lesion patterns that cut across existing behavioral measures. Moreover, the ability to disentangle regions with distinct effects on the same behavioral variable would offer a more nuanced understanding of brain-behavior relationships and improve the precision of clinical prediction and intervention.

Here we present a novel, data-driven approach to the identification of the lesion-behavior maps. Causal Feature Learning (CFL) is a machine learning method we developed [25, 26, 27] with three key desiderata in mind:

1. Discover causal relations between high-dimensional causes (here: voxel level lesion maps) and high-dimensional effects (here: multivariate behavioral measures on neuropsychological tests).
2. Do not fix the granularity of the cause or of the effect a priori. Instead, learn from the data how to align lesion types with behavioral patterns without specifying candidate types or patterns in advance.
3. Do not presuppose a functional form of the causal relation (such as e.g. linearity).

These desiderata are fundamental to a data-driven approach to understanding cognition. The first implies that causal considerations have to be integrated into the data analysis from the ground up to ensure that techniques for confounder control and estimation can be applied to the identified mapping. The second implies that it is an empirical question whether the lesion-behavior mapping is coarse or fine-grained, i.e. whether behavioral outcomes are sensitive to small differences in the lesions or whether larger differences in lesions are required for behavioral change. The third point accounts for the more complex interactions we can expect in the relation between lesions and behavior. Inevitably, the degree to which one can satisfy these desiderata will depend on the available sample size, the empirical distribution of samples and the granularity of the measures we start out with. But such “sample dependence” is entirely appropriate: With limited data, only coarse causal predictions about broad behavioral effects are supported; with dense sampling, we can resolve a finer-grained causal mapping between lesions and behavior. The key is that the framework for analysis should not hinge on restrictions to these desiderata.

### Causal Feature Learning

The motivation for Causal Feature Learning can be illustrated with a toy example developed by Spirtes & Scheines (2004)[28]: Suppose one is studying the effect of Total Cholesterol (*TC* ) on Heart Disease (*HD* ), but that *TC* is made up of two components, High Density Lipids (*HDL*), and Low Density Lipids (*LDL*), which, for the sake of argument, have different causal effects on *HD* (positive effect for *HDL* and negative for *LDL*) (Fig. 1). In that case the causal effect of *TC* on *HD* is necessarily ambiguous, because it depends on the proportion of *HDL* versus *LDL* that make up *TC*. With respect to its effect on *HD*, *TC* is an over-aggregated quantity and it would be misleading to describe it as a cause of *HD*. Instead, the correct level of causal granularity is that *HDL* and *LDL* are causes of *HD* (each with different effects). In contrast, if *HDL* and *LDL* had (quantitatively) the same causal effect on *HD* (say, improvement by three units), then they could be aggregated to *TC*, which would then constitute an appropriate cause of *HD*. The causal effect of *TC* on *HD* would then be unambiguous, or, definite. Definite effects needn’t be deterministic, as long as the probability of the outcome given the cause does not change with different instantiations of the cause, i.e. here the ratio of *HDL* to *LDL* makes no difference to the probability of *HD* given a value of *TC*.

**Figure 1:**
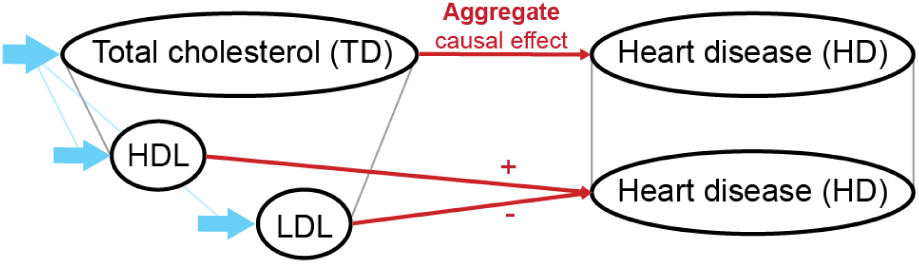
Toy example (adapted from [28]): Identifying the correct level of aggregation is crucial for causal inference. Suppose *HDL* and *LDL* have different causal effects on heart disease, but the researcher incorrectly aggregates them up to total cholesterol. In that case the causal effect of total cholesterol on heart disease will be ambiguous (even under intervention (blue arrows)), because it depends on the proportion of *HDL* vs. *LDL*. But if *HDL* and *LDL* had exactly the same effect on heart disease, then the aggregation to total cholesterol would be appropriate, as its causal effect on heart disease would be unambiguous (=definite). The Causal Feature Learning method identifies in a data-driven way the appropriate granularity of the causal relation by aggregating fine-grained effects while ensuring that the causal effect of the aggregate remains definite.

CFL is explicitly designed to preserve definite causal relations in the aggregation from the fine-grained measurement data to the causal representation, while not over-aggregating to causes that result in ambiguous effects. CFL does not assume that the granularity of either the cause or effect are pre-defined; instead, it discovers them from data. CFL takes paired high-dimensional variables (*X, Y* ) as input, where *X* represents the cause space (here: high-dimensional lesion data), and *Y* the effect space (here: the high-dimensional behavioral scores). The goal is to determine whether there is a lower-dimensional (coarser) representation (*C, E*) of (*X, Y* ) that still captures the causal relation between *X* and *Y* . In the present work the goal is to identify groups of lesions associated with specific groups of cognitive deficits, where both the cause and effect groupings need to be learned.

CFL is based on the graphical models account of the causal effect of *X* on *Y* in terms of the do-operator, *P* (*Y |do*(*X*)), which specifies the probability distribution of *Y* given an intervention on *X* that controls the values of *X*. If *X* and *Y* are not confounded (nor subject to selection bias), then *P* (*Y |do*(*X*)) is just equal to *P* (*Y |X*). Importantly, this account of the causal effect makes the causal effect of *X* on *Y* independent of the marginal distribution *P* (*X*) over the cause *X*, thereby capturing the fundamental idea of the independent mechanisms that underlie causal relations [29] and ensuring that the causal effect on *Y* is well-defined even if one (interventionally) changes the distribution over *X*.

CFL first estimates the high-dimensional conditional distribution of *P* (*Y |X*). It then coarsens *X* and *Y* while preserving the conditional probability. The goal is to identify the granularity of representation that provides the basis for a causal interpretation (for intervention and manipulation), not merely a predictive one. Formally, it groups *X* values *x*_1_ and *x*_2_ together if they have the same conditional distribution (“Cause aggregation”):

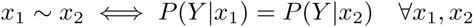

On the effect side, two outcomes *y*_1_ and *y*_2_ will be grouped if their conditional probabilities *P* (*y*_1_*|X*) and *P* (*y*_2_*|X*) have the same profile for any *X* = *x* (“Effect aggregation”):

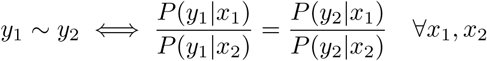

This condition ensures that any distinction that is preserved in the coarsening of the effect *Y* can be controlled by a change in the state of *X*. The result is a coarsening of the state spaces of *X* and *Y* into partitions whose cells correspond to variables *C* and *E*, respectively, that are sufficient statistics for *P* (*Y |X*). Learning *P* (*Y |X*) and clustering *X* and *Y* on the basis of this relation can be achieved in a variety of ways depending on data size and dimensionality.

If *X* and *Y* are confounded, the resulting *C* and *E* need not reflect causal features because *P* (*Y |X*) may differ from interventional probability *P* (*Y |do*(*X*)). However, the Causal Coarsening Theorem [25] shows that, under mild conditions, the partitions obtained from *P* (*Y |X*) will almost always be refinements of the causal partition. That is, *C* and *E* will contain all causally relevant distinctions but may also include non-causal ones due to confounding. There are then two paths forward: One is to experimentally test the distinctions in the observational partition to determine which are causal versus merely predictive (such as in [25]). Alternatively, if intervention is impossible (as it is here), one must test for confounding using observational data. Here, we consider a variety of measured demographic quantities as candidate confounders and check whether they correlate with the lesion categories CFL identifies. If so, we also check whether they correlate with the behavioral outcomes conditional on the lesion categories to establish confounding. We can then compare whether the dependence between lesion categories and behavioral outcomes is entirely accounted for by the measured confounder or whether there remains residual dependence. While this check addresses confounding by observed quantities, the possibility of further unmeasured confounding cannot be excluded without stronger assumptions.

We thus obtain a data-driven lesion-behavior map that links anatomical features (lesion location) to particular behavioral response patterns (questionnaire responses). CFL automatically identifies the optimal cause and effect granularities: lesions at possibly distinct locations that have the same effect on behavior, and behavioral responses that are related with respect to lesion effects. Conversely, lesions with different effects on behavior are separated *even if* they overlap in voxel space.

On the effect side, individuals with similar *aggregate* behavioral responses (e.g. similar total scores on questionnaires) may be distinguished from one another if the lesion data reveal item-level dissociations. In such cases new, possibly cross-cutting behavioral aggregates can be identified. Alternatively, when established behavioral measures track lesion effects, CFL provides an independent confirmation of those measures and suggests brain regions that are candidate causes of the behaviors these measures capture.

Like Canonical Correlation Analysis (CCA), CFL permits a brain-wide search for lesion-behavior relations without restricting analyses to small, region-defined subsamples (e.g., [30], which contrasted 7 ventromedial prefrontal cortex (vmPFC) with 5 dorsolateral prefrontal cortex (dlPFC) lesion samples). But unlike CCA, which compresses high-dimensional *X* and *Y* into components that maximize correlation, CFL disaggregates the distinct ways in which *X* can influence *Y* and imposes no parametric form on the lesion-behavior map (given sufficient sample size). Both CCA (at least in principle) and CFL permit multivariate outcomes, but – as shown in the next section – CCA’s representations are difficult to interpret relative to ground truth, while CFL straightforwardly describes both the cause and effect features and possible interactions within the cause space.

In the machine learning literature there are a variety of alternative approaches to learning causal representations. In particular, variational auto-encoders (VAEs) also provide non-parametric compression approaches. While limited studies have used VAEs to map lesions to behavior [31], constraining such models typically requires sample sizes that are generally not available in this domain (the dataset used in [31] is roughly 10 times the size of the dataset used here). Additionally, unless specifically designed to model causal systems, VAEs are essentially compression tools that do not provide any causal guarantees – nothing ensures that *X* will not be over-aggregated, as in the toy cholesterol example. However, when sufficient samples are available, these machine learning tools can substitute subroutines in CFL where we currently use less data hungry methods.

To illustrate the CFL analysis pipeline, we start by showing in a set of simulations, with known ground truth causal relations, that CFL (a) accurately recovers brain regions that cause a particular behavioral outcome, (b) correctly identifies and distinguishes lesions that have distinct effects on behavior, and (c) offers a natural causal representation and interpretation of the findings (in contrast to other methods). We then apply CFL to real lesion and behavioral data and recover well-known findings connecting left hemisphere lesions to language deficits and right hemisphere lesions to visuo-spatial deficits [32, 33]. Finally, we consider depressed mood, where lesion-symptom associations have been difficult to establish despite several decades of study [34]. It is possible that attempts to infer brain-behavior associations in depression have been challenging, in part, because depression is a syndrome that includes distinct types of symptoms that may individually localize to different anatomical systems. We apply CFL to Beck Depression Inventory (BDI) item-wise responses to recover regions associated with higher and lower ratings of depressive symptoms. We additionally test whether a total BDI summary score for depression provides the best measure for the lesion-depression symptom mapping given the available data, and how it compares to alternative ways of aggregating BDI questions. Our analysis suggests that the BDI summary score provides a good baseline to characterize this relation, but that a particular sub-grouping of the BDI questions discovered by using CFL provides a superior representation for relating lesions to depressive symptoms.

## Results

### Participants, behavioral measures, and neuroimaging data

All participants provided written informed consent and all procedures were approved by the Institutional Review Boards of the participating institutions and are in accordance with the Declaration of Helsinki. The patient cohorts for the following three analyses were drawn from two databases. 520 subjects were included from the Neurological Patient Registry of the Division of Neuropsychology and Cognitive Neuroscience at the University of Iowa Department of Neurology. The Registry includes patients who suffered brain damage from ischemic stroke, subarachnoid hemorrhage, or neurosurgical resection. Neuroimaging was done in the chronic epoch (> 3 months) after lesion onset. The second dataset included 195 individuals from the Vietnam Head Injury Study (VHIS) registry, which is composed of American veterans who suffered brain damage mainly from penetrating head injuries in the Vietnam War. Data collection details for all cohorts can be found in Methods: “Subjects.”

**Cohort 1:** Lesion masks from 455 individuals with unilateral lesions from the Iowa Registry. All lesions were represented on the left hemisphere, with right hemisphere lesions being flipped on the x-axis to increase spatial coverage for the simulation. A heatmap of lesions in this cohort is shown in Fig. 2A. Deficits were simulated as a linear function of lesion overlap with one of 100 left hemispheric regions from the Schaefer-200 parcellation, a commonly used representative parcellation [35] (see Methods: “Simulated behavioral deficits” for details and Fig.S10 for parcellation visualization). The distribution of an example deficit is shown in Fig. 2D.

**Figure 2:**
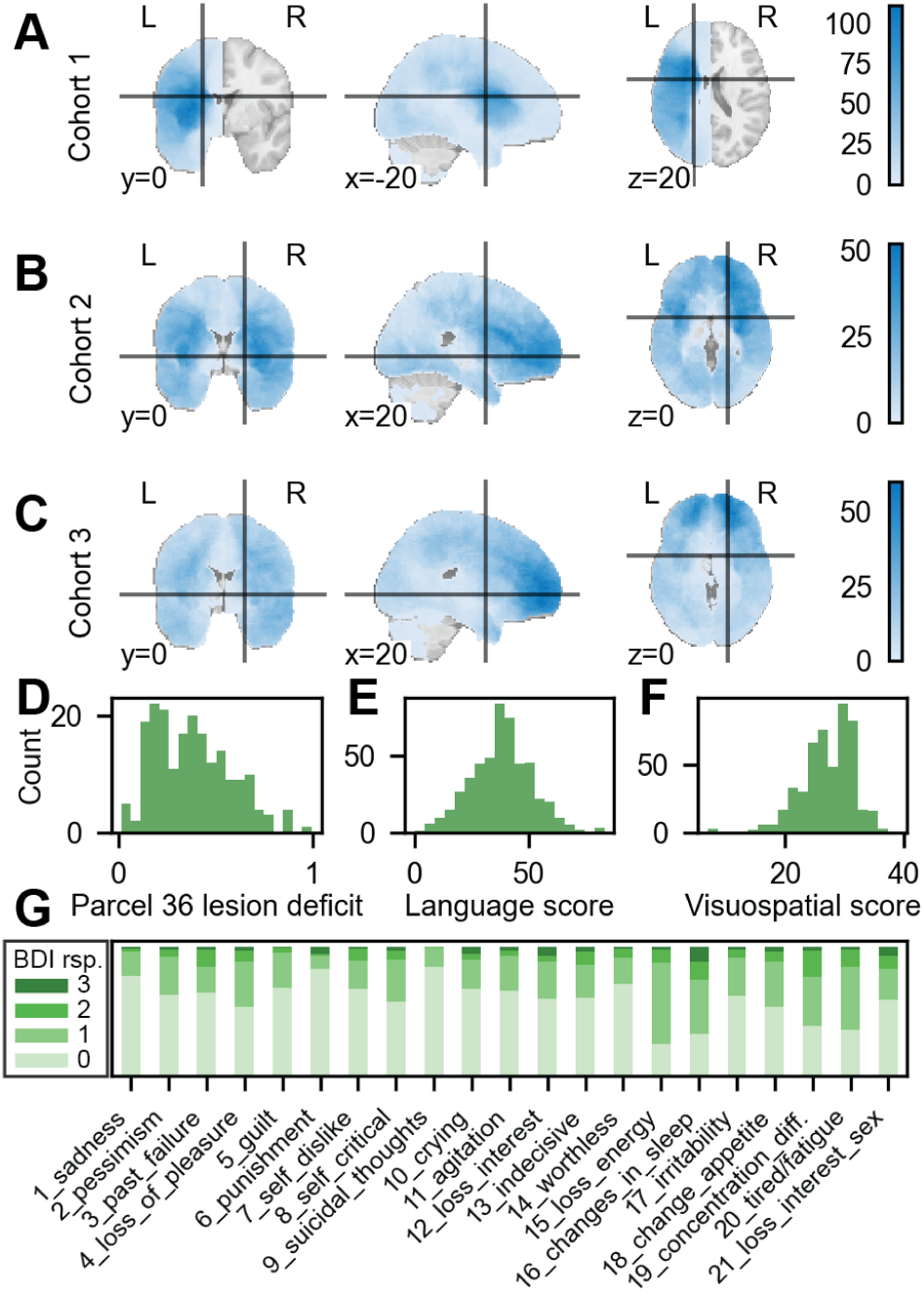
Cohort summaries. Heatmaps of lesion density across individuals for A) Cohort 1, B) Cohort 2, and C) Cohort 3 at central representative slices (heatmaps across all axial slices can be found in Fig. S1). Crosshairs indicate the locations of slices in other perspectives. Lesion size distribution can be found in Fig. S7. D) Distribution of an example simulated deficit caused by lesions to an example parcel. E) Histogram of cohort 2 scores on the Controlled Oral Word Association test (COWA). F) Histogram of cohort 2 scores on the Judgment of Line Orientation (JLO) test (the Pearson correlation between the two scores is r = 0.15). G) Response distribution for cohort 3 participants on the 21 items of the BDI questionnaire.

**Cohort 2:** 501 individuals from the Iowa Registry. Each patient completed a battery of neuropsychological tests, in particular, the Controlled Oral Word Association test (COWA), which evaluates language production, and the Judgment of Line Orientation test (JLO), which evaluates visuospatial discrimination. A heatmap of lesions in this cohort is shown in Fig. 2B. Histograms of the COWA and JLO responses are shown in Fig. 2EF.

**Cohort 3:** 520 individuals derived from two data sources: 325 patients from the Iowa registry and 195 individuals from the VHIS. For the latter, neuroimaging was obtained 36-39 years post-injury. Itemized Beck Depression Inventory-II (BDI-II) responses are available for all participants. This 21-item self-report questionnaire evaluates depressive attitudes and symptoms over the prior two weeks. Lesion heatmaps for this cohort are shown in Fig. 2C, and response histograms for all items are in 2G. Correlations among BDI items and between aggregate cognitive-affective and somatic sub-scores [36, 37, 30] are provided in Fig. S9.

### Lesion-behavior mapping

Lesion-behavior maps were learned using Causal Feature Learning (CFL). Given the comparatively small sample size, we configured CFL with a ridge regression that predicts test scores from lesion masks to approximate *P* (*Y |X*), and a KMeans clustering model to partition lesion masks based on this learned relation. We chose the ridge regression regularization parameter *α* based on cross-validated prediction error. The number of clusters *K* used by the KMeans model was selected by optimizing the recoverability of the conditional distribution from the resulting partition. Model configuration, fitting, and hyperparameter tuning details can be found in the methods section (Methods: “Causal Feature Learning (CFL)”).

For each analysis, CFL categorized lesion masks according to the effects the lesions had on neuropsychological test scores. Lesions with similar causal effects are placed in the same category. For the BDI analysis, CFL additionally identified groups of itemized BDI responses that are similarly affected by the distribution of lesion masks. We compared this grouping to that which arises when we solely use the standard summary BDI statistic (sum of responses to BDI questions) to obtain an empirical evaluation of the suitability of this coarser summary statistic to describe depressive symptoms.

### Recovering simulated behavioral deficits with CFL

The first simulation ensures that in cases of known ground truth, CFL reliably recovers the correct lesion causing the deficit, and that it does so more accurately than CCA as the current established method for this task. Using Cohort 1, we compared CFL and CCA in recovering 100 ‘ground truth’ lesion-behavior maps, with each behavioral ‘deficit’ generated based on overlap with one of 100 brain regions. Evaluating the methods on standard lesion recovery measures, we find that CFL recovers ground truth regions with comparable or improved accuracy relative to CCA (Fig. 3; one-tailed Wilcoxon test; LESYMAP vs. CFL (within-category agreement threshold = 80%), CFL (90%), and CFL (95%), respectively; dice score: *p* = 0.009, 0.333, 0.995; peak displacement: *p* = 8.270e*−*7, 8.270e*−*7, 8.270e*−*7; centroid displacement: *p* = 0.005, 0.006, 0.004; contour displacement: *p* = 0.247, 4.684e*−*4, 1.284e*−*5).

**Figure 3:**
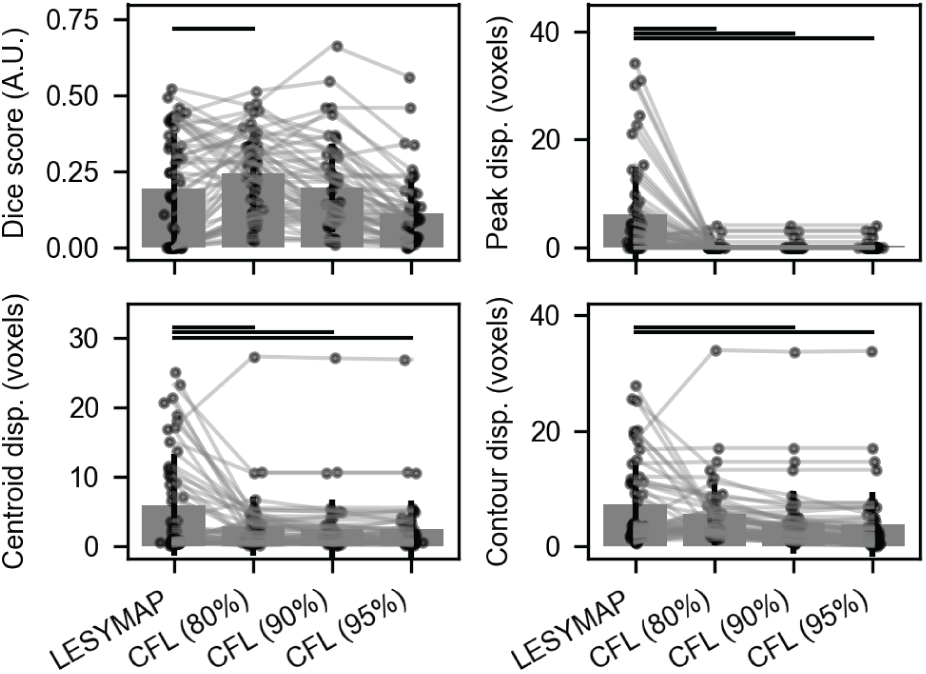
Comparison of CFL and CCA (LESYMAP) single-region lesion-behavior mapping. Lesion-behavior mapping performance by CFL and CCA (horizontal bars at top indicate *p <* 0.05, one-tailed Wilcoxon test between pairs of mapping approaches). Connected points correspond to a single deficit-generating region. CFL suffixes denote the level of within-category agreement required per voxel in addition to permutation testing (see Methods: “CFL: Deficit significance testing”). Higher value corresponds to better performance for dice score; lower value corresponds to better performance for all other scores.

Next, we compared CFL and CCA in the multivariate-multivariate setting. Using Cohort 1, we designated three of the 100 left hemispheric regions from the Schaeffer-200 parcellation, simulated distinct deficits for each, and computed the lesion overlap for each subject with all three regions simultaneously to then test how well CFL and CCA are able to disentangle multiple lesion-behavior mappings. See Fig. S11 for the ground truth regions used to generate the simulated behavioral deficit scores, associated deficits, and deficit correlations. We fit CCA with a three-dimensional bottleneck, capped by the number of deficits. Doing so yields weights mapping voxels to the three modes (Fig. 4A) and weights mapping deficits to the three modes (Fig. 4B).

**Figure 4:**
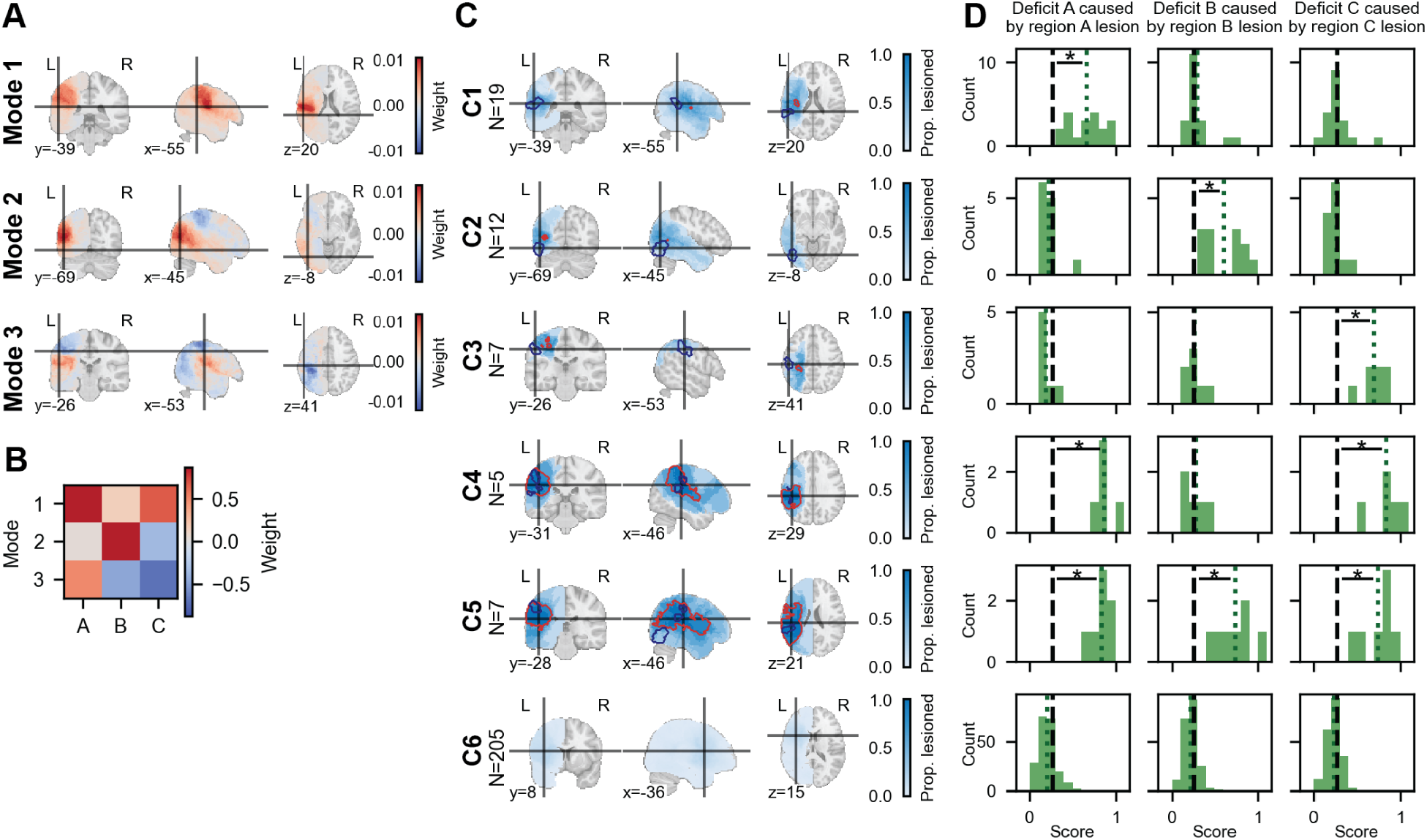
CFL maps lesions to behavior in an interpretable manner. A) CCA lesion weights, projecting lesion masks to three modes that maximize correlation with deficit modes. Color indicates the contribution of each voxel to each CCA mode. Crosshairs indicate the center of the largest contiguous region of significant voxels (outlined in red). These cross-sections were shown as opposed to those of the ground truth regions since the three CCA modes do not correspond to any specific ground truth region. B) CCA deficit weights, projecting deficit scores as a result of lesions to three regions A, B, and C to three CCA modes that maximize correlation with lesion modes. Color indicates the contribution of each deficit score to each CCA mode. C) CFL lesion categories and D) corresponding deficit distributions for deficits caused by lesions to regions A, B, and C. For panel C, color indicates the proportion of category members with a given voxel lesioned. Categorization is shown on withheld samples. For panel D, green dotted lines indicate observed means. Black dashed lines indicate mean of resampled population means (100,000 resamples, sample size equal to category size). Asterisk (*) indicates that the observed mean is significantly greater than the resampled means, *p <* 0.05, 100,000 resamples. Significance tested for categories 1-5.

In the multivariate-multivariate case, CCA identifies regions (and differences between regions in modes 2 and 3) that are associated with weighted combinations of the three deficits, instead of disambiguating the three distinct deficit-generating regions. This illustrates one of the problems with CCA: CCA modes are directions that maximize correlation, which do not necessarily correspond to distinct lesion-behavior relationships. In contrast, CFL identified six lesion categories that capture both the isolated effects of lesions on particular deficits, as well as interaction effects of lesions that lead to multiple deficits (Fig. 4C, see Fig. S2 for more detail). CFL-learned categories 1-3 isolate the three deficit-generating regions (associated deficit distributions compared against 100,000 permutations, *p <* 0.05, see Methods: “CFL: Deficit significance testing” for more details). Category 4 identifies lesions that affect both deficits A and C, and category 5 identifies lesions that affect all three included deficits. Category 6 also appropriately separates out those subjects with lesions that result in no or minor deficits. By partitioning the space of lesions based on the multi-dimensional probability distribution of the three deficits conditioned on lesions, CFL captures the lesion-behavior relations present in the data in a straightforward manner.

Finally, to highlight the importance of incorporating the lesion-behavior relation into learning from the outset, rather than first reducing lesion-space dimensionality, we ran KMeans directly on the Cohort 1 lesion population, using the Davies-Bouldin score to select the number of clusters (see Fig. S5). Aside from a common “catch-all” cluster, there is little agreement in the partitions found by these two approaches. KMeans clusters are driven solely by lesion location and size, grouping lesions with different behavioral deficits and separating lesions with similar deficits. This shows that coarsening before accounting for lesion-behavior relations discards critical information.

### CFL identifies established language and visuospatial brain regions from real behavioral data

To demonstrate that CFL can recover field-established results in a data-driven manner, we applied CFL to Cohort 2 to analyze the relationship between lesions and language and visuospatial performance. As was done in the previous analysis, for this analysis CFL was configured to map multivariate lesion masks to multivariate outcomes (a language production test score and a visuospatial discrimination test score). CFL identified five categories of lesions with distinct behavioral outcomes (Fig. 5, see also Fig. S3 and S6AC for more detail). Although CFL does not enforce spatial localization or contiguity, four of five categories show spatial specificity. Lesion category locations align with prior findings. Left-hemispheric lesions in categories C2 and C4 correspond to below-average language scores, relative to average Cohort 2 scores. The two categories distinguish between larger (C2) and smaller (C4) left-hemispheric lesions that correspond to more and less severe language effects, respectively (Fig. 5B). Permutation testing (see Methods: “CFL: Deficit significance testing” for details) determined language scores to be significantly lower for C2 and C4 (indicated by the asterisk). Right-hemispheric lesions in category C1 correspond to significantly below-average visuospatial scores, while language performance remains unaffected.

**Figure 5:**
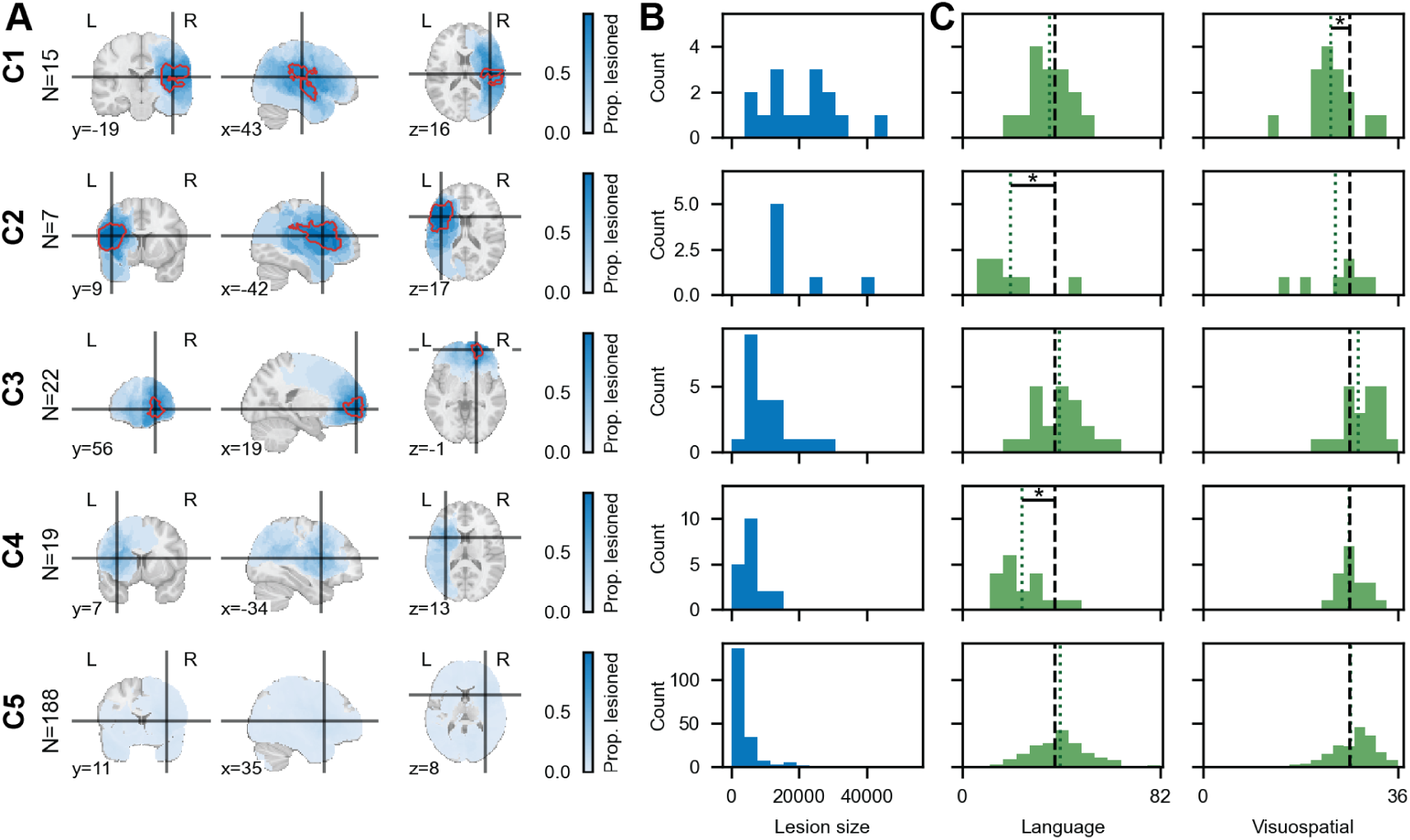
CFL finds five lesion categories with distinct outcomes on language and visuospatial tasks. A) Normalized lesion heatmap within each category at central coronal, sagittal, and axial slices. Color indicates the proportion of category members with a given voxel lesioned. Crosshairs indicate the locations of slices in other perspectives. Center of mass MNI coordinates: C1: (43*, −*19, 16), parietal operculum and posterior insula; C2: (*−*42, 9, 17), left inferior frontal gyrus of Broca’s area and underlying white matter; C3: (19, 56*, −*1), right frontal pole; C4: (*−*34, 7, 13), dorsal anterior insula and frontal operculum; C5: (35, 11, 8). Heatmaps across all axial slices can be found in Fig. S3. Red boundaries delineate significant voxels (see Methods: “CFL: Identifying significant voxels” for details). B) Lesion size (number of voxels lesioned) distributions associated with individuals in each lesion category. C) Distributions of normalized COWA (left) and JLO (right) scores associated with individuals in each lesion category. Green dotted lines indicate observed means. Black dashed lines indicate mean of resampled population means (100,000 resamples, sample size equal to category size). Asterisk (*) indicates that the observed mean is significantly different from the resampled means, *p <* 0.05, 100,000 resamples.

Prefontal cortex lesions in category C3 correspond to slightly above-average language and visuospatial scores, though these do not reach significance. It is important to interpret these results in the context of the sample - the majority of this cohort is patients with left and right middle cerebral artery (MCA) strokes, which decreases the average language and visuospatial scores for the overall sample. In contrast, C3 contains a larger proportion of patients with lesions due to surgical resection, which are intentional lesions designed to minimize deficits (see Fig. S8 for complete etiology distributions by CFL lesion category). Without baseline, per-patient measures of language and visuospatial performance prior to lesion onset, we cannot conclude that C3 implies enhanced performance as opposed to successful surgery.

Category C5, which includes the largest proportion of subjects, shows no lesion localization and corresponds to average language and visuospatial scores – lesions that don’t have specific effects on either test. This catch-all category is analogous to category C6 that we found in the simulations (see Fig. 4) that does not permit further differentiation given the samples and sample size we have.

The unsupervised delineation of left-hemispheric and right-hemispheric lesions and their respective associations with language and visuospatial deficits demonstrates the efficacy of CFL as a data-driven method for lesion-behavior mapping. The isolation of frontal cortex lesions corresponding to slightly higher language and visuospatial scores demonstrates the advantage of using a data-driven approach to identify all lesion-behavior relationships present in a dataset at the appropriate level of granularity. If we had instead taken a hypothesis-driven approach by comparing all left and right hemisphere lesions, then (1) finer-grained distinctions between lesions within each hemisphere would have been obscured and (2) the distinct (and opposite!) effect of frontal cortex lesions, which may occur in the left or right hemisphere, would not have been identified.

### Lesion categories are stable with respect to participant demographics

While CFL preserves causal information present in the fine-grained measurement data, it cannot rule out confounding effects without further assumptions or analysis. While the lesion itself is an intervention, lesion type may correlate with other factors that also directly affect behavior, thereby confounding the lesion-behavior relation. The most obvious candidates for confounding are demographic features that make particular lesions more likely while also directly affecting behavioral outcomes (i.e. their effect is not mediated by the lesion). We examined potential confounding due to available demographic measures: *age at time of lesion scan*, *sex*, *whether-Hispanic*, *race*, and *education*. *Whether-Hispanic* and *race* responses were homogenous across the cohort. To control for the remaining three demographic measures, we checked whether the demographic quantities correlated with the CFL-derived lesion categories – a necessary condition for confounding. We found no significant dependence between any of the demographic information and the lesion categories (Supp. Fig. S12A). Overall, these results are consistent with prior findings that the effects of lesions on cognitive abilities far outweigh any confounding due to demographic quantities [32].

We note that the lesion etiology mentioned as a candidate explanation for the above-average performance of subjects with prefrontal cortex lesions (C3 above) does not constitute confounding: We observe a positive association between prefrontal cortex lesions and behavioral outcome measures within Cohort 2 of lesion subjects, and most of the subjects in C3 have surgical resections. But that just illustrates the successful causal effect of precision surgery. It is not a case of confounding, since confounding would require a causal effect of the decision to have surgery on the behavioral outcome measures that is *not* mediated by the lesion. In fact, the observed dependence is fully accounted for by the lesion: for a given lesion category, we find that etiology does not further distinguish behavioral scores (to the extent this can be checked given limited etiology coverage in the dataset, S13).

Our analysis, however, does not preclude the possibility that the relation between lesions and cognitive scores is still confounded by some further *unmeasured* quantity. Without stronger assumptions or experimental intervention this possibility always remains, but here the interventional nature of the lesions helps: A confounder would have to cause the particular lesion type *and* also directly influence task performance.

### CFL identifies three lesion categories with distinct effects on BDI

Having demonstrated CFL on well-established language and visuospatial results, we next examined depression, an area with few established findings. With Cohort 3, we analyzed the relationship between lesion location and responses to 21 individual BDI items. Depression is a construct with distinct processes assessed via overall severity (total BDI score) or sub-scores (e.g. somatic vs. cognitive-affective). Beyond aggregating lesions, here we also sought the optimal granularity for the *effect* measure by grouping BDI responses with similar likelihoods given lesion location (see Methods: “Multivariate-multivariate CFL” for implementation details).

As before, the high-dimensional expected conditional probability of BDI responses given lesion location was estimated using ridge regression, and KMeans was used to partition the space of lesion masks. Effect coarsening was implemented with a second KMeans model applied to the estimated probability of each sample in the effect space given each cause category found in the previous algorithmic step. All free parameters were tuned in an unsupervised manner; details can be found in the Methods section.

CFL identified three distinct lesion categories and four distinct BDI response patterns (Fig. 6, see Methods: “Selecting number of categories” and Fig. S6BD for details). Cause category C1 identifies lesions localized in vmPFC, while C2 identifies dlPFC and insula lesions. Cause category C3 contains non-localized lesions that result in all effect categories as expected under the null distribution, with the exception of E2, which is unlikely, but slightly more common than expected in this population (see Methods: “CFL: Significance testing causal relation” for details). Even though category C1 identifies specific regions, its associated BDI scores largely match those of the catch-all C3 category. In contrast, C2 is strongly associated with high BDI scores (lower than expected loading on E1 and E2 and strong loading on E4). Note that this dataset did not include any bilateral dlPFC lesions, which limits the maximum voxel-wise agreement within these regions in C2.

**Figure 6:**
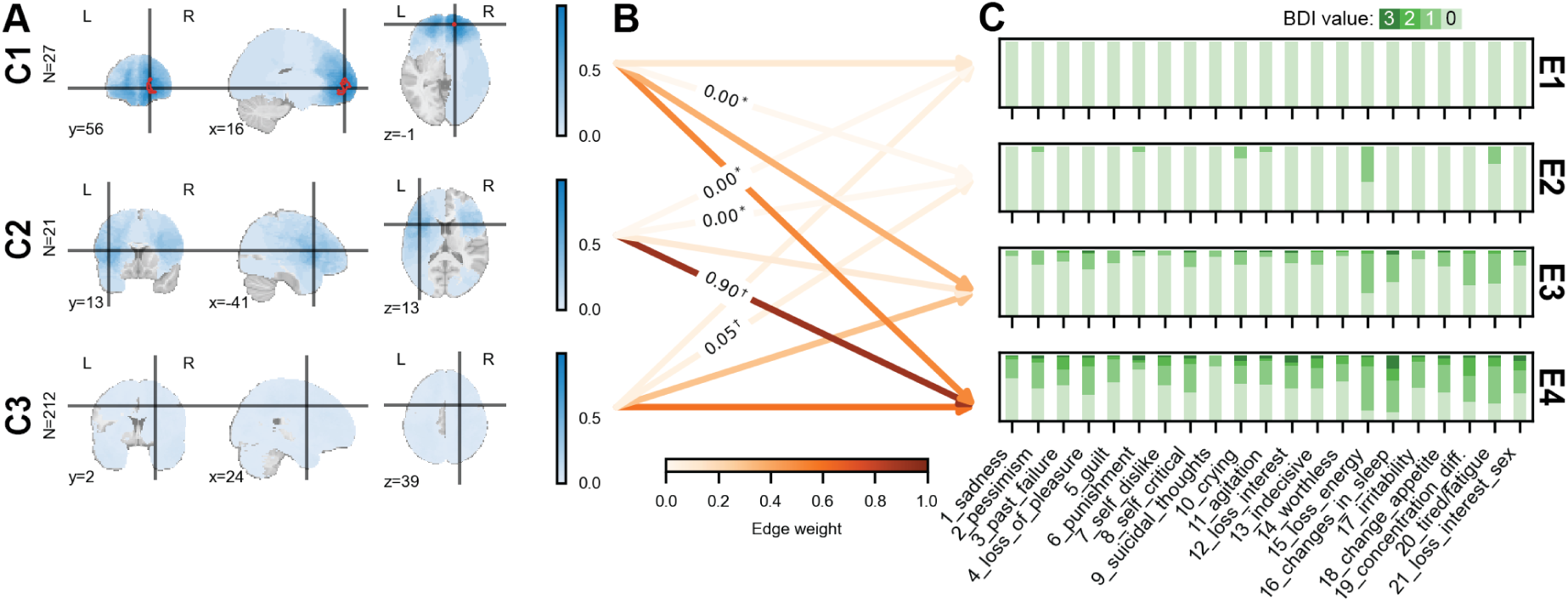
CFL maps vmPFC lesions to milder BDI scores and dlPFC lesions to more severe BDI scores. (A) Normalized lesion heatmap within each category at central coronal, sagittal, and axial slices held constant across categories. Crosshairs indicate the locations of slices in other perspectives. Color indicates the proportion of category members with a given voxel lesioned. Center of mass MNI coordinates: C1: (16, 56*, −*1), C2: (*−*41, 13, 13), C3: (24, 2, 39). Red boundaries delineate significant voxels (see Methods: “CFL: Identifying significant voxels” for details). Heatmaps across all axial slices can be found in Fig. S4. (B) Probability of each BDI effect category given each lesion cause category. Only edges that are significantly greater (*†*) or less than (*) expected under the null hypothesis (see Methods: “CFL: Significance testing causal relation”) are annotated (note that this may be the case even for very weak edges in the case that the edge weight is significantly less than expected). Note that all possible edges are drawn, even if the edge has value 0, in order to visualize significance. C) Distributions of responses to each of the 21 BDI questions within each BDI effect category identified by CFL. Categories are sorted from best (E1) to worst (E2) overall outcomes. (See Methods: “Partition of behavior space” for details.)

These findings are notably distinct from past studies and those we presented in Fig. 5 in that those related particular lesion profiles to unidimensional distributions of JLO and COWA scores. While the scores shown in Fig. 5 are the commonly used summary scores, here in Fig. 6 we are aiming to distinguish unique profiles of item-wise BDI scores. As was done with the language-visuospatial dataset, a demographic analysis showed no evidence of confounding by age, sex, and education. As Supplementary Figure S12B shows, there is no significant dependence between demographic quantities and the identified lesion categories.

Identifying vmPFC and dlPFC/insula as regions of “resilience” and “risk” with respect to depression is consistent with the findings of previous hypothesis-driven analyses that compare the two regions exclusively [30]. In line with more recent data-driven efforts to relate lesion masks to a unidimensional summary BDI score using LESYMAP [38], the CFL analysis shown here provides strong additional support for these findings by considering lesions throughout the brain in a large sample. The analysis here furthers our ability to understand the relationship between lesions and depressive symptoms in its capacity to simultaneously account for both multivariate lesion mask and multivariate behavioral scores in an interpretable manner. As will be shown in the next section, this framework offers a novel opportunity to determine whether additional relations exist between lesion categories and more fine-grained BDI subscores that may have been obscured when considering BDI as a single summary score.

### Interpretation of depression effect categories found by CFL: Five BDI summary variables are sufficient to recover CFL lesion categories

The preceding analyses considered the 21 questions on the BDI individually. However, BDI is often summarized as a one-dimensional total or mean BDI score. With a comprehensive categorization of lesions based on the full 21-dimensional score, we can now ask whether the mean BDI score is sufficient to identify this categorization or whether collapsing the 21 BDI questions down to one score discards information crucial to the lesion-behavior relation.

To answer this question, we re-ran CFL with the mean BDI as the target variable and compared the resulting lesion categories to the three categories identified above with the 21 questions as the target variable (Fig. 6). We found that the lesion categories based on the mean BDI score can recover the original partition to some extent, but that C3 lesions are often miscategorized as C1 and C2 (Fig. 7B). We performed the same analysis on another commonly used set of three summary variables for BDI – the averages of somatic (orange in Fig 7A), cognitive-affective (blue in Fig 7A), and non-loading scores, respectively [36]. We found that this representation of BDI reduced the number of misclassifications, but that C3 lesions were still misclassified as C1 and C2 (Fig. 7C).

**Figure 7:**
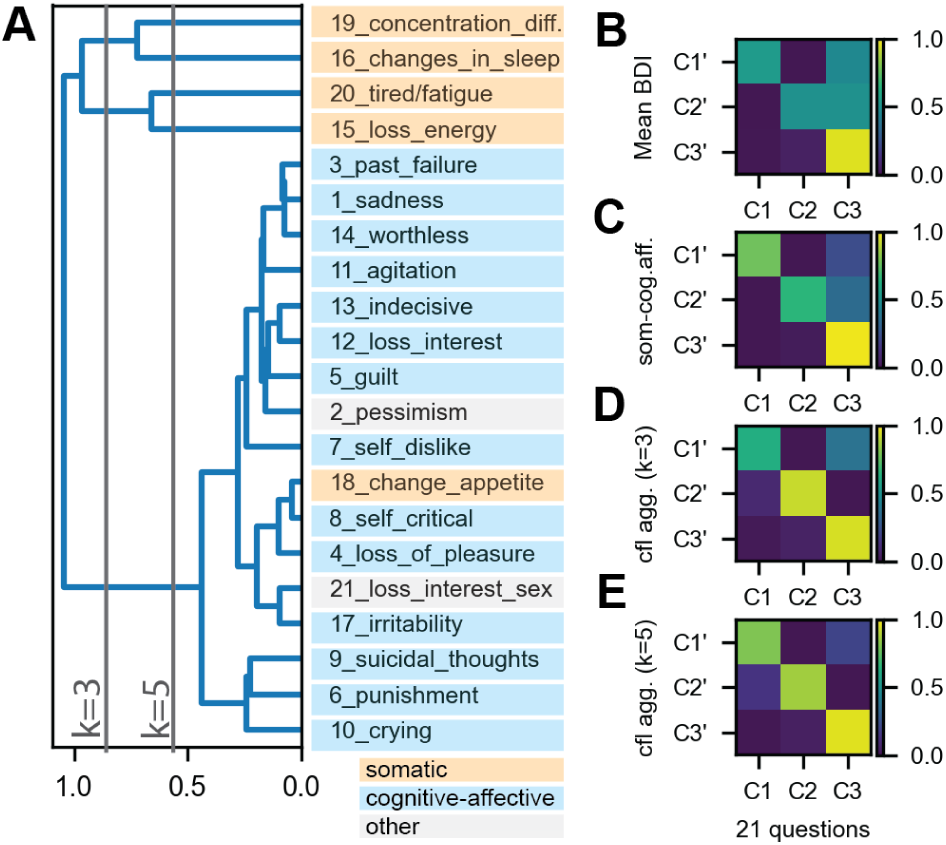
Clustering 21 questions by importance in defining CFL lesion category boundaries recovers somatic vs. cognitive-affective distinction, and further refines each grouping. A) Hierarchical clustering of 21 BDI questions based on similarity in dissociating effect categories. B) Comparison in ability to recover CFL lesion categories using mean BDI. C) Same as B but with somatic and cognitive-affective aggregates. D) Same as B but with 3 question clusters found by CFL (see k=3 bar in A). E) Same as B but with 5 question clusters found by CFL (see k=5 bar in A).

To identify which distinct groups of questions are needed to largely recover the lesion categories found with all 21 questions, we designed a similarity measure across questions based on the extent to which they each discriminated between pairs of CFL effect categories (see Methods: “Constructing BDI summary variables with CFL” for details). We used this similarity measure to cluster questions in a hierarchical manner (Fig. 7A). We then asked what degree of clustering was necessary to correctly classify the C3 lesions that were misclassified using the aforementioned BDI summary variables. We found that when fitting CFL with averages of *k* = 3 question clusters, C3 samples misclassified as C2 were largely eliminated but that C3 samples misclassified as C1 remained (Fig. 7D). *k* = 5 question clusters were needed to minimize C1 misclassifications (Fig. 7E). This clustering amounts to averaging across cognitive-affective questions and treating four of the five somatic questions independently: 19: concentration difficulty, 16: changes in sleep, 20: tired/fatigue, and 15: loss of energy. Together, these findings suggest that while cognitive-affective questions can largely be summarized by a single average variable, treating somatic questions independently (with the exception of change in appetite) – and not as a single aggregate quantity – is necessary to separate out the lesion-depressive symptom relations we found with the full 21 question representation.

Overall, this analysis provides a case study of how to simultaneously aggregate both lesions and the behavioral measures to a granularity that optimally describes their relation. The inferred granularity is, of course, subject to the limitations of the availability of varied fine-grained data, both for the lesions and for the 21 questions. However, given these limitations, the analysis empirically confirms that the standard BDI score as a simple sum of the itemized responses offers a good separation of the different lesion categories and depression outcomes. This is somewhat improved by the combination of cognitive and somatic measures, but the best separation is achieved by an outcome measure that further subdivides the somatic questions.

## Discussion

### Summary

We applied a new data-driven representation learning method, Causal Feature Learning (CFL), to a set of three related lesion-behavior mapping questions. First, we showed that CFL accurately maps lesion location in relation to simulated behavioral deficits with known ground-truth associations. The discovered lesion categories were more accurate and more directly interpretable than the results obtained from Canonical Correlation Analysis. They also differed starkly from the lesion clusters found by naive clustering on lesion anatomy only. Second, CFL recovered the expected relationships between left hemisphere lesions and language deficits and right hemisphere lesions and visuospatial deficits using real behavioral data [32, 33, 39, 40, 41]. These results remained stable under checks for potential confounding due to measured demographic quantities.

Having demonstrated that CFL can recover well-established lesion-behavior relationships with simulated and real-world behavioral data, we turned to the unresolved relationship between lesions and depressive symptoms. CFL found dlPFC and insula lesions to disproportionately relate to moderate/severe depressive symptoms, and vmPFC lesions to relate to the distribution of BDI symptoms expected under the null distribution. Upon comparing BDI response categories that CFL identified with the conventional summary statistics for BDI, we found that the mean BDI score adequately summarizes the information about the effects of lesions on behavior that is captured by the BDI, but that a superior representation can be achieved with a finer splitting of the somatic questions that further isolates tiredness-related questions.

Importantly, the relationships found in both analyses were identified in an unsupervised manner with minimal investigator input. Given a dataset of lesions throughout the brain and the associated full distribution of cognitive test scores, CFL produced these well-established relationships without guidance. The fact that CFL teased apart the dichotomies between left and right hemispheric lesions in relation to language/visuospatial ability, and between vmPFC and dlPFC in relation to BDI [30], lends data-driven support for past work that has recovered the same relationships. It also demonstrates how future lesion-behavior mapping analyses can be accelerated by replacing hypothesis-by data-driven methods that infer the relevant brain regions. Furthermore, the analysis of the BDI effect space demonstrates how CFL can be used to relate lesions to fine-grained behavioral data across individual questionnaires and even across broader assessments of behavior that span multiple domains of function, offering a route to finding new cross-cutting lesion-behavior relations. Few methods offer a data-driven approach to identifying the appropriate granularity to analyze a lesion behavior map, and CFL does so while ensuring that all the standard tools to establish causal claims (e.g. to check for confounding) can still be applied. A method for predicting task performance for new lesion subjects is intrinsic to the learned mapping, and to the extent that the absence of confounding is established, the learned lesion categories can provide guidance on how to improve clinical intervention.

### Limitations

Though lesion studies are much closer to interventional than most human neural data, the observed lesion and measured behavioral performance may still be confounded by unmeasured quantities. We have attempted to control for confounding due to measured demographic quantities, but further unmeasured confounders might have led CFL to form additional distinctions between categories that do not reflect the causal relationship of lesions on behavior. For instance, particular lesion distributions are systematically associated with particular etiologies, such as temporal lobe lesions with epilepsy. And epilepsy subjects may (independent of the lesion) also exhibit particular task performance patterns, resulting in a correlation between lesions and task performance that is not a causal effect of the lesion. While epilepsy subjects were excluded from this analysis, this caveat still applies to similar sources of unmeasured confounds and must be kept in mind when interpreting results.

Relatedly, given the uncontrolled nature of natural lesion occurrences, the results are biased by the locations of the brain that have been well “sampled” by the lesions. The lesions in our sample of patients did not cover all areas of the brain; lesions varied in size and particular anatomical areas will have been lesioned more often than others. As a result, even though there might be sharply delineated causes of behavior in the brain, CFL may not be able to precisely aggregate to the correct granularity given the available data, and some ambiguity about the optimal granularity can result. Both larger sample sizes and efforts to enrich samples to ensure the most isotropic anatomical distribution could help with these limitations. Another note is that certain lesions are systematically excluded – namely, those that result in impairments thought to produce invalid task performances (e.g., blindness would be an exclusionary criterion for performing the JLO), so the effects of those brain regions cannot be appropriately assessed.

A final limitation lies in our ability to visualize and interpret the discovered categories, even once the above limitations have been acknowledged. Though visualizations of average lesion masks and lesion volume distributions allow us to compare lesion location and volume across categories, these methods may not capture other lesion characteristics that may be driving the causal relations that CFL identifies. Further work in visualizing the more subtle lesion characteristics that may constitute the crucial causal distinctions between the identified categories would allow for a more complete understanding of CFL results. A particularly important class of these would be distinctions between damage to gray matter and white matter, where the latter would require further interpretations in terms of disconnection of brain areas as opposed to the absence of particular brain areas.

### Future work

As noted, nearly all of the lesions in our study include white matter and can thus be thought of as producing disconnection syndromes, at least in part. This challenge also points to the broadest recommendation for the future: combine methods. Applying functional neuroimaging to lesion patients, and combining lesion analysis with white matter atlases and/or diffusion-weighted imaging could all contribute to producing an essential level of description that provides the interface between the lesion masks we obtained and the behavioral impairments with which they are causally associated [42]. Given that CFL is agnostic to the type of input and target data to relate, the method demonstrated in this work can naturally be extended to include not only lesioned regions but also disconnections between regions as input features to relate to behavioral measures. That level of analysis would help describe the mechanism whereby the lesion causes the change in behavior: through a distributed change in brain function. Such altered brain function could reflect not only impairments in cognition but also compensation or even improvements [43, 44].

CFL provides an opportunity to systematically evaluate these brain-behavior relationships in a principled way. Just as diverse measurements of brain activity could be integrated on the cause side, a broad set of human behaviors spanning cognition, personality, and affect could be included on the effect side. CFL could then find anatomically informed relationships across these behavioral dimensions, providing both an empircal test of existing findings and a data-driven means to uncover novel relations that may cut across traditional accounts of brain-behavior relations.

## Methods

### Subjects

Participants included 585 individuals from the Patient Registry of the Division of Neuropsychology and Cognitive Neuroscience at the University of Iowa Department of Neurology who met study criteria. The Registry includes patients who suffered brain damage from ischemic stroke, subarachnoid hemorrhage, or neurosurgical resection. Neuroimaging was done in the chronic epoch (> 3 months) after lesion onset. Inclusion criteria for this study included the presence of a stable brain lesion. Each major analysis (simulation, language–visuospatial performance, depressive symptoms) used a distinct cohort defined by available behavioral data. The simulation analyses (cohort 1) used lesion masks from 455 individuals; only unilateral cases were included, and right-hemisphere lesions (identified by visual inspection) were flipped to the left to increase spatial coverage. Cohorts 2 and 3 comprised individuals who completed the relevant behavioral assessments, including neuropsychological testing and the Beck Depression Inventory-II (BDI-II) [45]. Exclusion criteria included a neurological or psychiatric disorder that preceded the brain lesion, or a history of significant alcohol or substance abuse. Cohort 3 additionally included 195 subjects drawn from the W.F. Caveness Vietnam Head Injury Study (VHIS) registry, which consists of military veterans who suffered penetrating head trauma while in combat during the Vietnam War era (1967–1970). Inclusion criteria for this study included individuals with lesion structural neuroimaging and BDI-II completion in the chronic phase postinjury, defined as > 3 months later. All VHIS combat veterans had been declared fit for duty at the time of their enlistment in the military prior to their head injury. In accordance with federal and institutional guidelines, all participants from both registries provided written informed consent, and all procedures were approved by the University of Iowa Institutional Review Board and are in accordance with the Declaration of Helsinki.

### Simulated Behavioral Deficits

For Cohort 1, deficits were simulated as a linear function of lesion overlap with one of 100 left hemispheric regions from the Schaefer-200 parcellation [35] (Fig. S10A) plus Gaussian noise (Fig. 2D). For ground-truth brain region *R_i_*, the corresponding deficit *D_i,j_*for lesion mask *L_j_*was calculated as:

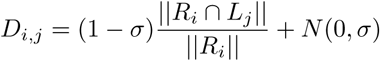

where *σ* = 0.1 and *|| ∗ ||* is the number of voxels. Deficits for each ground truth region were then scaled to a range of [0, 1] across subjects. Higher scores correspond to larger simulated deficits. Deficit-generating regions varied in lesion size (Fig. S10B).

### Cognitive Assessment

Participants in the Iowa Registry performed a large battery of cognitive tests (see [45] for details of individual tests). We used the Controlled Oral Word Association (COWA) task of verbal fluency from the Multilingual Aphasia Examination (MAE) with additive corrections by age and education (score range before corrections: 0 to the number of words the subject can generate in one minute), and the Benton Judgement of Line Orientation with additive corrections by age and sex (30 questions, score range before corrections: 0-30). COWA and JLO scores were independently z-scored prior to analysis.

### Mood Assessment

Mood was assessed using the Beck Depression Inventory-II (BDI-II) [36], a 21-item self-reported questionnaire that evaluates characteristic attitudes and symptoms of depression experienced over the preceding two weeks. Items correspond to affective, cognitive, somatic, and vegetative symptoms of depression that align with the criteria used to diagnose major depression in the fourth edition of the Diagnostic and Statistical Manual (DSM-IV). If participants had repeat assessments, the highest score was selected. BDI scores were collectively z-scored prior to analysis.

### Lesion Segmentation

Each included subject had a focal brain lesion with visible boundaries evident from research-quality structural imaging from T1 and T2 sequences on MRI. CT was used in rare cases in the Iowa Registry when MRI was contraindicated (*n* = 75 of 585 Iowa subjects across both analyses). Imaging for all subjects was performed in the chronic epoch (> 3 months since onset). The anatomical segmentation of lesion borders was traced manually for each subject and brought to a common template space for statistical analyses. The MAP-3 method of lesion tracing involves the manual tracing of lesion borders on a template brain using the lesion depicted in an MRI or CT scan as a guide and has been previously described [46, 47]. With improvements in automated methods for transforming brains to a common space, lesions traced after 2006 were manually traced on native T1-weighted scans with FSL [48] and then transformed to the 2mm MNI152 atlas using nonlinear registration and lesion masking techniques available in ANTs [49]. Because lesions negatively affect the accuracy of the transformation to MNI space, transformations for unilateral lesions were performed using enantiomorphic normalization [50]. The approach replaces the lesion volume with the voxel intensities from its non-damaged homolog to more closely align the individual’s brain with the template brain. Bilateral lesions were transformed by applying a cost function mask to the lesion volume [51], which reduces the influence of voxels within the lesion volume on the transformation process. The spatial transforms were applied to the brain and lesion mask. The anatomical accuracy of the lesion tracing was reviewed in native and MNI space and edited as needed by a neurologist (A.D.B.) blinded to mood ratings.

Lesion masks were taken at 2*mm*^3^ voxel resolution (lesion volume ranges: cohort 1: 48*mm*^3^*−*754, 704*mm*^3^; cohort 2: 120*mm*^3^ *−* 430, 200*mm*^3^; cohort 3: 120*mm*^3^ *−* 781, 784*mm*^3^; full distributions in S9. Lesion mask volumes were vectorized for all analyses. Only voxels located within the brain volume (as opposed to all voxels in the three-dimensional rectangular prism that encapsulates the brain volume) were included.

### Causal Feature Learning (CFL)

While CFL applies across data domains, the account here is tailored to lesion–behavior mapping and holds for all analyses unless noted. With the exception of the first cohort 1 CFL analysis, which performs multivariate-univariate mapping for each of 100 simulated deficits, all CFL analyses are performed as multivariate-multivariate mappings. For a general, detailed account of CFL theory and implementation, see Chalupka et al., (2016) [52]. A full CFL implementation can be found at [27].

Letting *X* be vectorized lesion masks and *Y* be multi-dimensional cognitive assessments, the goal of CFL is to maximally coarsen *X* and *Y* to a granularity that still preserves the causal relationships present in the micro-level data. To do so, CFL groups together micro-level values that play the same (or very similar) roles in the relationship between *X* and *Y* .

First, CFL estimates the conditional probability *P* (*Y |X*) from data. Since density estimation is challenging in high dimensions, we approximate this step by estimating the conditional expectation of the distribution E[*P* (*Y |X*)], as was done in [52]. This amounts to regressing *Y* on *X* with the mean-squared-error measure.

#### Conditional probability estimation

For all analyses, we estimated E[*P* (*Y |X*)] with ridge regression. Regularization hyperparameter *α* was chosen through 10-fold cross-validation over log-spaced values. The value of *α* that minimized out-of-sample error was chosen. The following values of *α* were used for cohorts 1,2, and 3 analyses, respectively: 1*e*3, 4.6*e*4, 1*e*6.

A single layer neural network produced qualitatively similar results but showed instability across initializations given the limited sample size for the high-dimensional lesion masks.

#### Partition of lesion space

Given a dataset (*x, y*) *∈ D*, CFL uses the estimate for E[*P* (*Y |X* = *x*)] *∀x ∈ D* to partition the space of lesion masks *X* . To preserve the relationship between *X* and *Y*, CFL groups points in *X* together that give rise to the same distribution in *Y*. In practice, we use KMeans to cluster samples *x*_1_*, x*_2_ *∈ D* together if E[*P* (*Y |X* = *x*_1_)] *^∼^*= E[*P* (*Y |X* = *x*_2_)] (a relaxation of the equivalence relation defined in [52]). Of course, equality in expectation does not ensure equality in distribution, and can be particularly misleading when the outcomes are mixtures. It is an approach we adopted from [52] to cope with the low sample size in the high-dimensional space.

#### Selecting number of categories

To choose the number of clusters *K* for KMeans in an unsupervised manner, KMeans was fit for varying values of *K*, with random initialization and 50 restarts. For each cluster assignment, we measured how much information was retained about the conditional probabilities that were clustered (Fig. S6AB). We quantified this by taking the least-squares estimate of the conditional probabilities from one-hot-encoded cluster assignments and computing a 10-fold out-of-sample mean-squared error. We selected the largest value of *K* that still resulted in a meaningful decrease in this error, at which point the bulk of the relational information had been captured by the clustering. In each case we checked whether neighboring values of *K* resulted in categories with significant differences in their lesion distributions (Fig. S6).

#### Partition of behavior space

The following methods only apply to the Cohort 3 analysis. To coarsen the behavior space *Y*, CFL groups samples in *Y* together if they have the same likelihood of arising given any value in *X* . In other words, samples *y*_1_*, y*_2_ *∈ D* are grouped together if

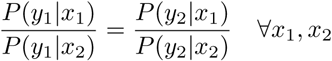

To compute the likelihood distribution at a given value of *Y* in practice, we take advantage of the fact that all *x ∈ X* that are in the same cause category have similar conditional probabilities by construction. Therefore, we only need to approximate the likelihood of each *y ∈ D* given each cause category. For each sample *y*, we approximate this likelihood with the average distance to its *K*-nearest neighbors in each cause category [52].

This approximation method relies on the idea that samples from the distribution will be closer to each other in regions where the density is high, relative to where the density is low. For each sample *y*, the calculated distances per cause category are normalized so that they sum to 1. These likelihood estimates were clustered using KMeans and the aforementioned hyperparameter tuning method to select the value of *K* to partition the effect space.

In summary, the CFL pipeline returns two pieces of information: it assigns each sample in our dataset to (1) a cause category (in this case, a lesion category) and (2) an effect category (for cohort 3, a depression category).

Specifically, in the analysis of Cohort 3, we estimated the probability of each 21-dimensional BDI-response *y_i_* in our sample given each of the three lesion categories identified. We then clustered two such responses *y*_1_ and *y*_2_ into one category *E_j_* if their conditional probability satisfied the equation above. The result is a partition of the high-dimensional effect space as shown in Figure 6C. Here the 21-dimensional space of questionnaire responses is partitioned into four effect categories (E1-E4), which we illustrate by showing the distributions of response patterns to the 21 questions within each category.

### Cohort 1 analysis: multivariate-univariate CFL and CCA

We ran CFL and CCA on the simulated Cohort 1 dataset described above. For each of the 100 simulated deficits, we fit CFL and CCA on a training dataset of 200 lesions to recover the associated deficit-generating region. CFL was configured as described in Methods: “Causal Feature Learning (CFL)”. The LESYMAP implementation of CCA was used with the following parameters: *method=“sccan”, optimizeSparseness=True, pThreshold=0.05 (default), smooth=0.4 (default), minSubjectPerVoxel=10 (default)*. Hyperparameters (ridge regularization scale for CFL, sparsity constraint for CCA) were fit using k-fold cross-validation within the training dataset. For CFL, the number of clusters for KMeans was fixed to 2 for all regions to maximize the contrast for the deficit area, though choosing the number of clusters in a cross-validated manner for example subjects found *K* = 2 as well.

With the model fixed, 255 test dataset lesions were assigned to the two categories by CFL. Of the two lesion categories, that which corresponded to a higher average deficit score was selected as the recovered deficit-generating region. For CFL, significant voxels were determined as described in Methods: “CFL: Deficit significance testing”).

For each model and deficit, we evaluated the recovered deficit-generating region *B* against ground truth region *A* using the following established metrics (Fig. 3E).

*Dice score:* a measure of spatial overlap, given by 2(*A*&*B*)*/*(*||A||* + *||B||*, where *|| ∗ ||* denotes the number of voxels.

*Peak displacement:* 0 if the peak of B falls within A, otherwise distance from the peak of B to closest voxel in A. For CFL, the peak corresponds to the voxel that had the maximum number of subjects who had that voxel lesioned. For CCA, the peak corresponds to the maximum of the SCCAN statistic computed by LESYMAP.

*Centroid displacement:* same as peak displacement, but with centroid of B instead of peak.

*Contour displacement:* average distance from contour of B to the closest voxel in A.

Performance metrics are only included for regions for which both methods yielded significant results (Table 1).

**Table 1:** Number of ROIs with significant results, by method.

| Model | # ROIs with significant results |
| --- | --- |
| LESYMAP | 61/100 |
| CFL (80%) | 84/100 |
| CFL (90%) | 82/100 |
| CFL (95%) | 74/100 |

### CFL: Identifying significant voxels

Voxels that (A) were lesioned more often than expected under the null hypothesis of no association between CFL category and voxel lesion rate and (B) had high within-category agreement on lesion status (at least 80% of subjects with a given voxel lesioned; see Fig. 3E for different percentage thresholds of inclusion) were included in the final recovered regions. To determine criteria (A), a permutation test was performed by shuffling CFL category labels 10,000 times. For a category of size *k* in the observed population, this process yields 10,000 resampled *k*-sized populations. For said category, a voxel was considered statistically significant if lesioned more often in the observed population category than in the corresponding *k*-sized category in at least 95% of the resampled populations.

### Cohort 1 analysis: multivariate-multivariate CFL and CCA

We ran CFL and CCA with the Cohort 1 dataset, including simulated deficits generated from three of the 100 originally defined regions. Unlike the previous Cohort 1 analysis, all three deficits were provided to CFL simultaneously. For CCA, the top 30 principal components (explained variance = 79%) of the lesion data were included, as is typically done with CCA when feature dimensionality is high [24]. As LESYMAP only supports multivariate-univariate mapping, the scikit-learn implementation of CCA was used instead with the three deficits included simultaneously. No additional constraints were applied. CCA was fit with a bottleneck dimensionality of 3, constrained by the deficit dimensionality. Both models were fitted to 200 samples. CFL hyperparameters were selected using cross-validation within the training set. Models were tested on 255 withheld test samples.

### CFL: Deficit significance testing

CFL lesion category labels were permuted 100,000 times. For each lesion category of size *k*, this provided 100,000 *k*-sized deficit samples. For each category, the mean of each resampled population was compared to the mean of the observed sample to determine if the observed sample differed significantly from a random population of the same size. If the observed deficit population was greater than (or less than, depending on the analysis) at least 95% (Bonferroni corrected when multiple deficit measures were tested) of resampled populations, it was considered statistically significant.

### CFL: Testing potential confounders

Candidate confounders (age at time of lesion scan, biological sex, and years of education) were tested for dependence with the CFL-identified lesion categories using a Two-sided Mann-Whitney U test comparing category-specific to full cohort distributions. Absence of a significant dependence between candidate confounder and lesion category indicates absence of confounding due to that candidate confounder. Presence of a dependence would necessitate further checks for a dependence between candidate confounder and behavioral outcome that is not mediated by the lesion categories. But since none of the available candidate confounders here passed the first criterion, this step was never needed.

For the case of lesion etiology, which did vary with lesion category in Cohort 2, dependence between etiology and the behavioral outcome was tested as follows. Conditioning on each lesion category, the distributions of test scores per etiology were compared to one another to determine whether etiology provides additional information about the behavior beyond what lesion category already provides. For each pair of etiologies (with more than five samples for the given lesion category in the dataset), the difference in test score distributions was quantified with a two-sided t-test to confirm no significant variation in test scores as a function of etiology.

### CFL: Significance testing causal relation

To test the significance of relations between cause and effect categories (Fig. 6B), the association between cause and effect category labels was shuffled 100,000 times. On each shuffle, the cause-effect transition matrix was recomputed. If the observed probability of transitioning from a given cause category to a given effect category was greater or less than that in 95% (Bonferroni-corrected) of shuffled samples, it was considered significantly greater or less than expected under the null hypothesis, respectively.

### Constructing BDI summary variables with CFL

CFL returns a partition over all possible response patterns in the 21-dimensional space of BDI question, but such a partition gives little insight into what questions are doing the work to define partition boundaries. To come up with such an organization over the questions, we grouped questions together based on how important each question was to define the state space boundaries that CFL constructed.

For each BDI question *q_i_*, we computed an “importance vector,” where each entry corresponds to a pair of effect categories *M_a_*and *M_b_*, and its value is the KL-divergence between the distribution of *q_i_* values in *M_a_* and *M_b_*. This importance vector tells us how distinct *q_i_* responses are between pairs of categories – the larger the distance between *q_i_* distributions in *M_a_* and *q_i_* in *M_b_*, the more information *q_i_* has about whether an effect sample should be in category *M_a_*or *M_b_*.

We hierarchically clustered (centroid linkage method, Euclidean distance) the 21 questions by their importance vectors, grouping questions that similarly discriminate between BDI categories. The mean responses for each question group provided “CFL aggregate” quantities. We varied the number of question clusters to find that which largely recovered the partition found with all 21 questions when CFL was rerun with the aggregate quantities.

### Alternative BDI summary variables

*Mean BDI:* all 21 questions were averaged. *Somatic vs. cognitive-affective:* Questions were averaged according to the groupings determined in [30]. Somatic questions: loss of energy, changes in sleep, changes in appetite, concentration difficulty, tiredness/fatigue. Cognitive-affective questions: sadness, past failure, loss of pleasure, guilt, punishment, self-dislike, self-critical, suicidal thoughts, crying, agitation, loss of interest, indecisiveness, worthlessness, irritability. Uncategorized questions: pessimism, loss of interest in sex.

### Software versions

python=3.8.20; scikit-learn=1.3.0; nilearn=0.10.4; LESYMAP=0.0.0.9222

## Author contributions

I.W. & F.E. performed the data analysis and led the writing of the paper. J.Griffis, R.A. & A.B. contributed to analysis and writing. J.Grafman, A.B. & D.T. collected and processed the data.

## Funding acknowledgments

F.D.E. and R.A. were supported by the Chen Neuroscience Institute and the Carver Mead New Adventures Fund. A.D.B. and J.G. were supported by NIH-NINDS 5R01NS114405, and the Roy J. Carver Charitable Trust. F.D.E., R.A., and I.A.W. were supported by the Schmidt Academy for Software Engineering fellows program. F.D.E. was supported by NSF EAGER (BCS-1845958).

## Supplementary Figures

**Figure S1:**
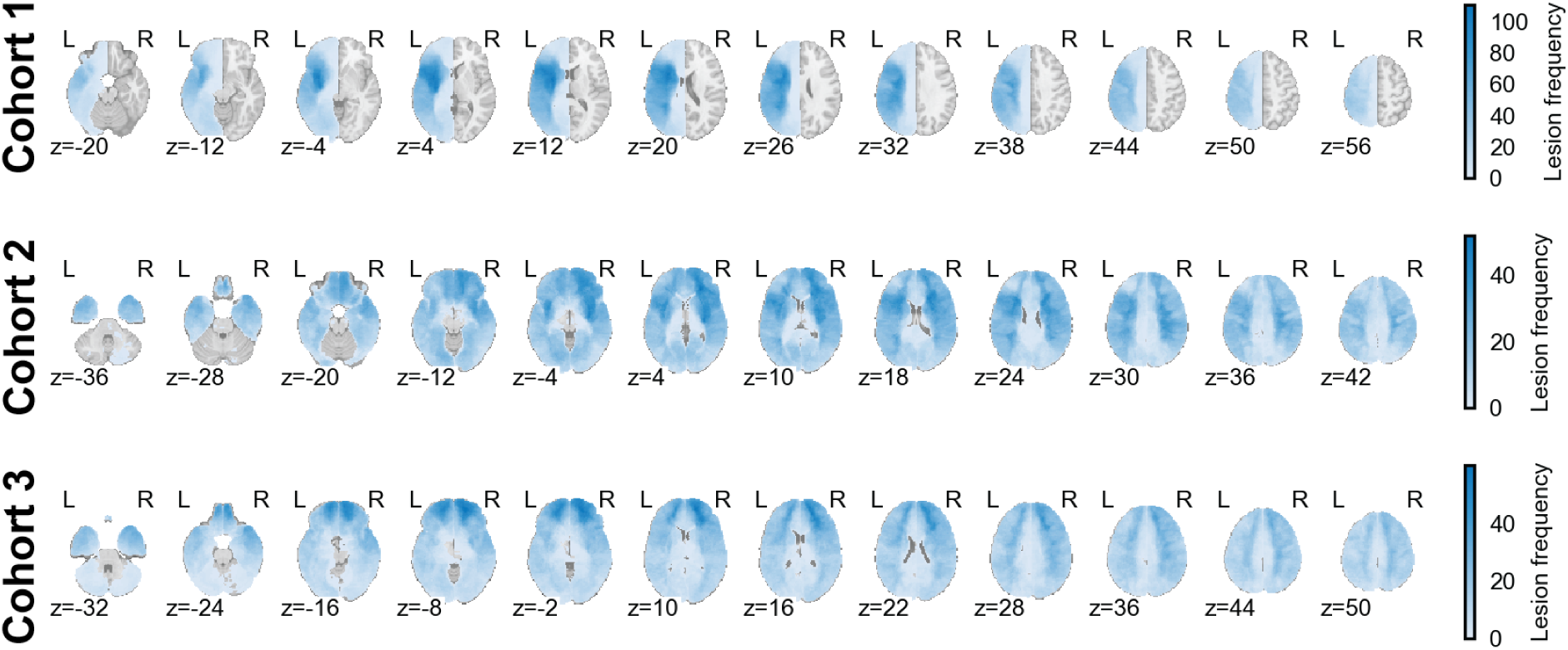
Heatmaps of lesion density across individuals for Cohort 1 (top), Cohort 2 (middle), and Cohort 3 (bottom). Axial slices are shown at evenly spaced z-coordinates. Color indicates the number of individuals in the cohort with a given voxel lesioned. See Figure 2 for more details.

**Figure S2:**
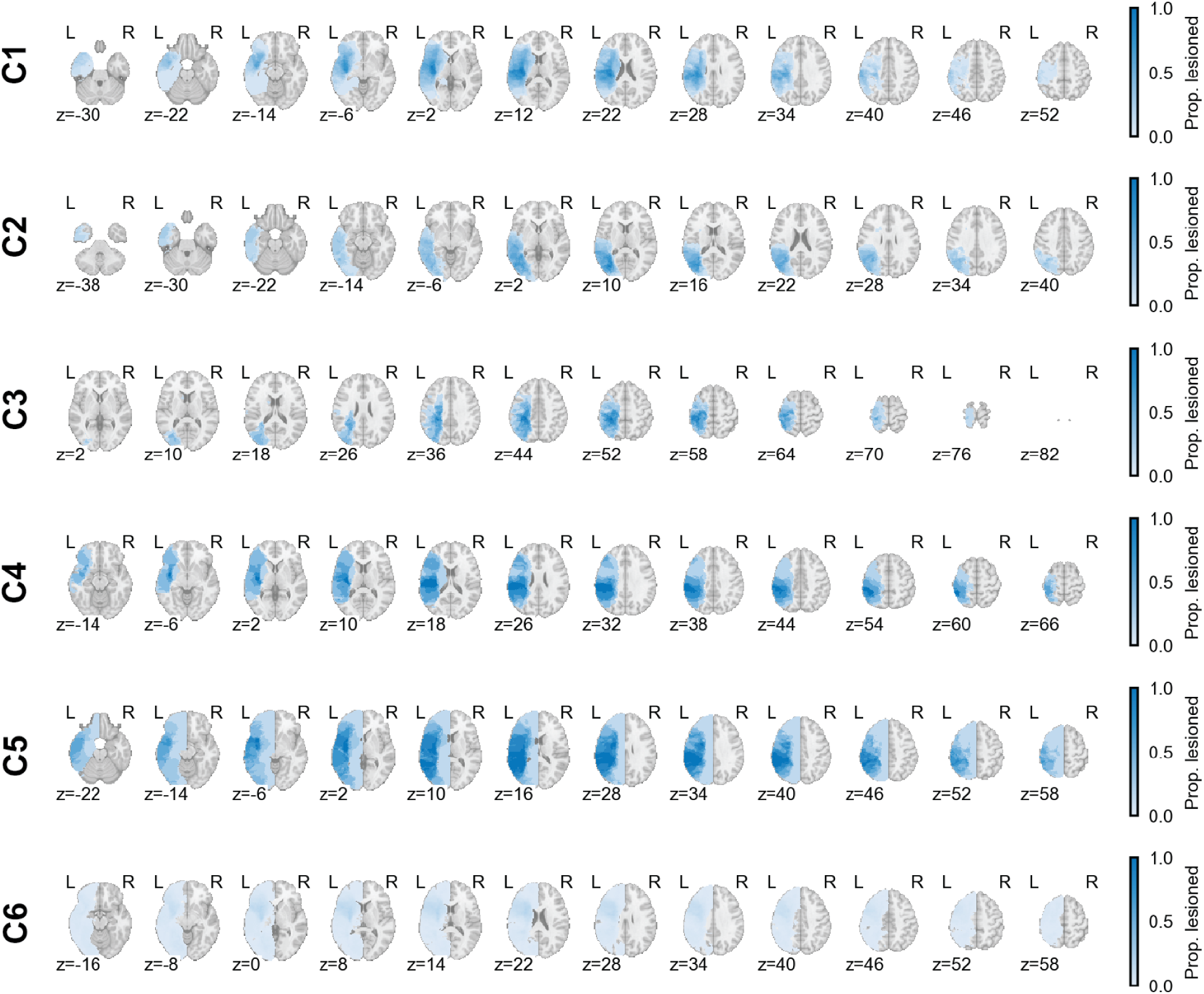
Heatmaps of lesion density across individuals in each CFL macrostate for the simulated Cohort 1 analysis. Axial slices are shown at evenly spaced z-coordinates. Color indicates the proportion of individuals in a macrostate with a given voxel lesioned. See Figure 4 for more details.

**Figure S3:**
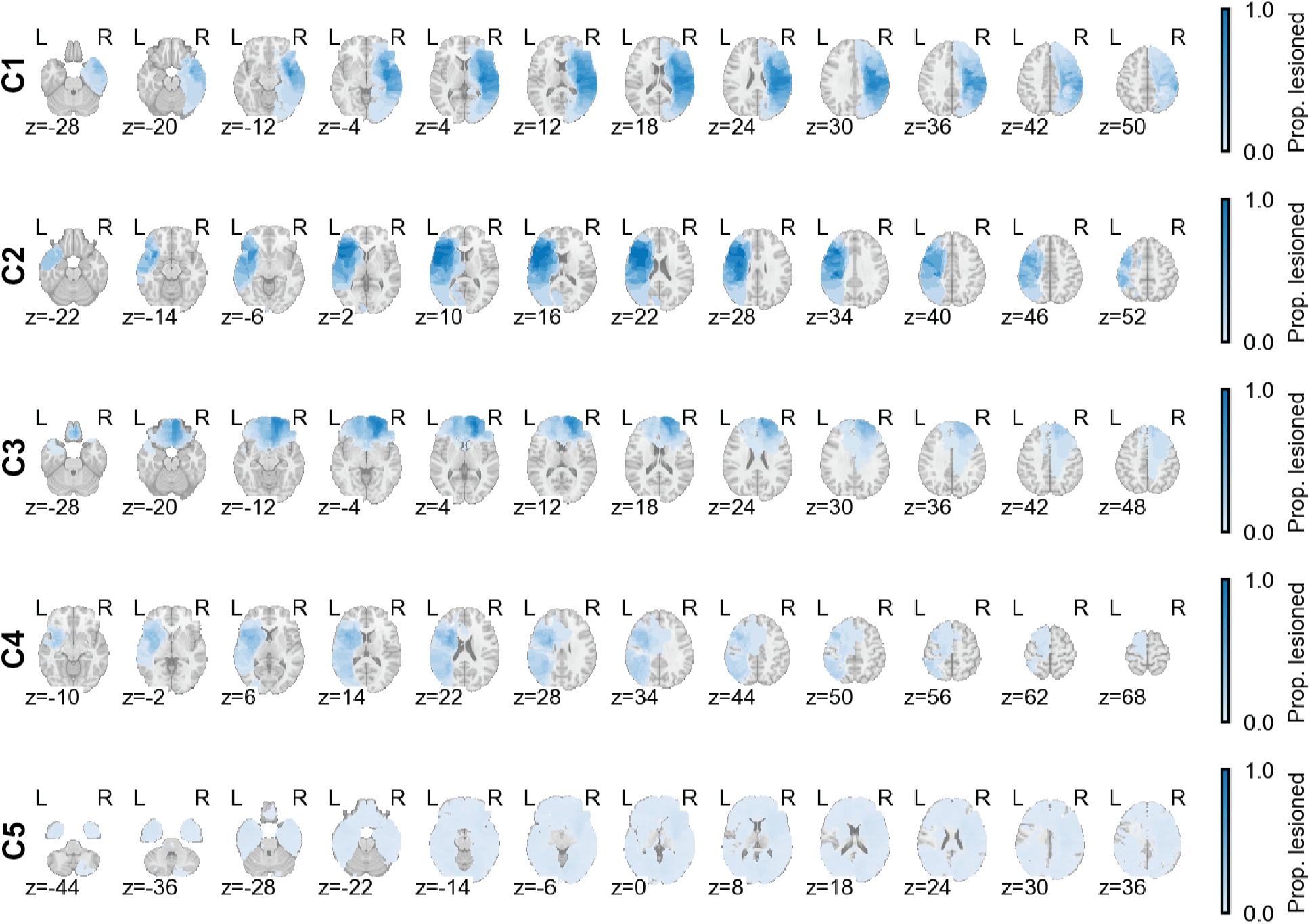
Heatmaps of lesion density across individuals in each CFL macrostate for the language-visuospatial analysis (Cohort 2). Axial slices are shown at evenly spaced z-coordinates. Color indicates the proportion of individuals in a macrostate with a given voxel lesioned. See Figure 5 for more details.

**Figure S4:**
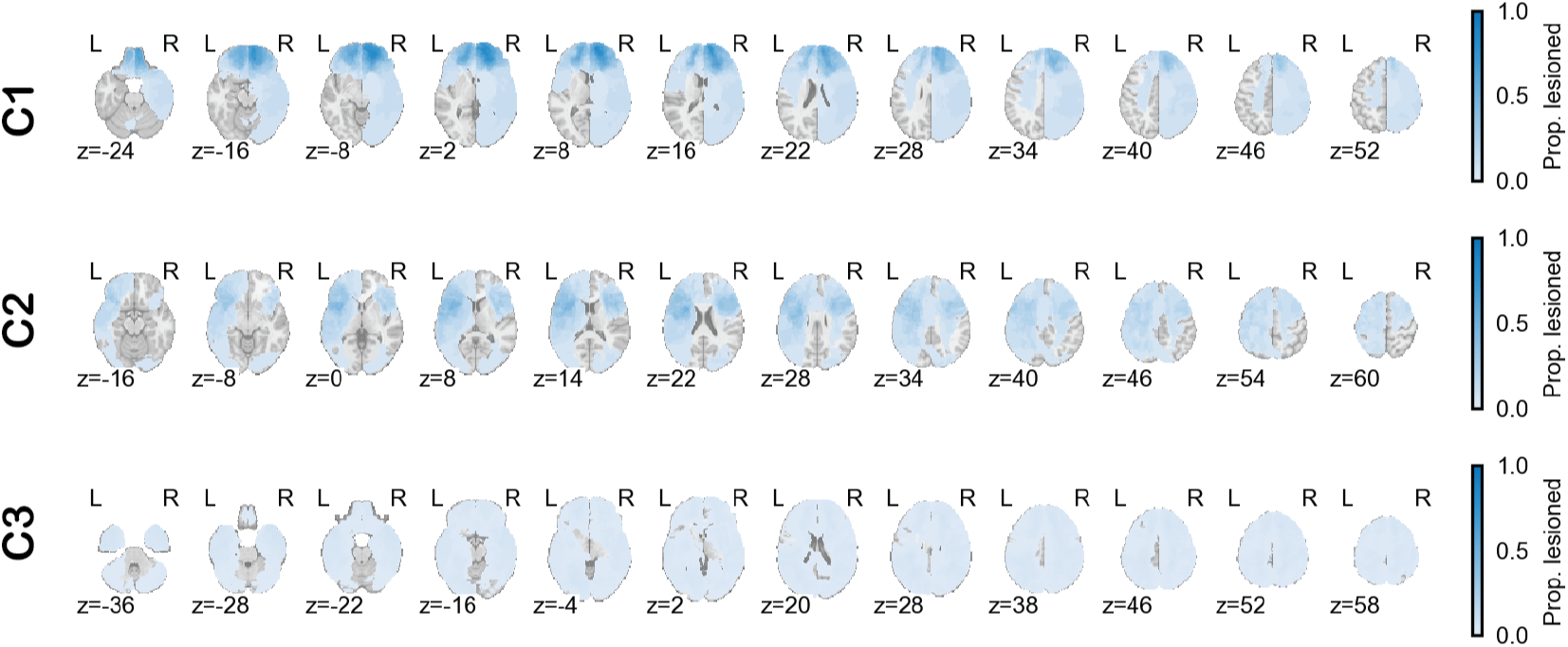
Heatmaps of lesion density across individuals in each CFL macrostate for BDI analysis (Cohort 3). Axial slices are shown at evenly spaced z-coordinates. Color indicates the proportion of individuals in a macrostate with a given voxel lesioned. See Figure 6 for more details.

**Figure S5:**
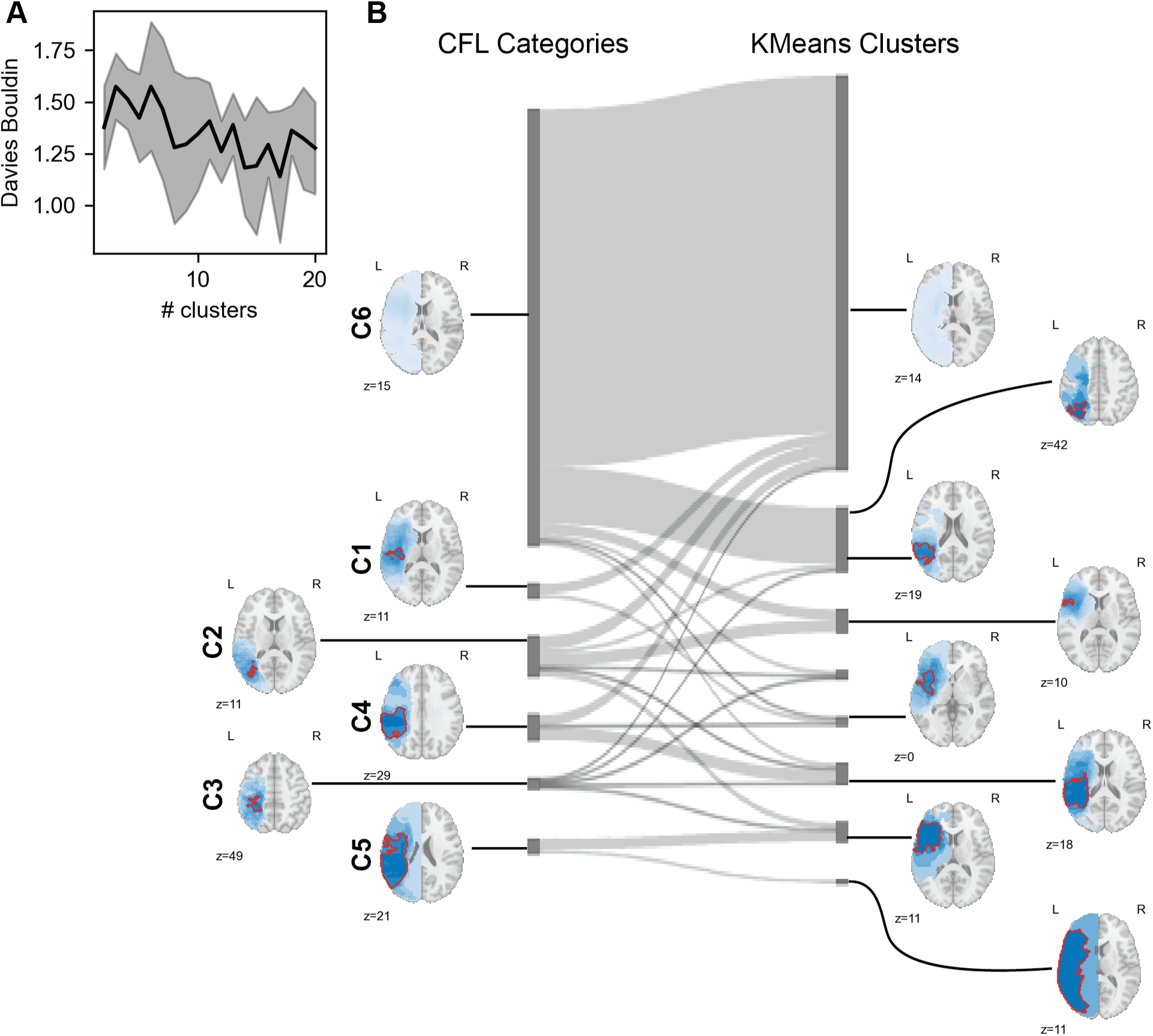
Lesion mask clusters found by KMeans applied only to the lesions cut across CFL categories that were learned from the lesion-behavior relation. A) Davies-Bouldin goodness of clustering score as a function of number of clusters found by KMeans. There is no obvious ideal cluster number, but *k* = 8 marks the first significant drop. B) Sankey diagram comparing cluster membership identified by CFL (left) and KMeans (right). Thumbnails show lesion mask category/cluster averages. Red boundaries delineate significant voxels (see Methods: “CFL: Identifying significant voxels” for details). Thumbnails are shown at a slice aligned with the center of the largest contiguous region of significant voxels.

**Figure S6:**
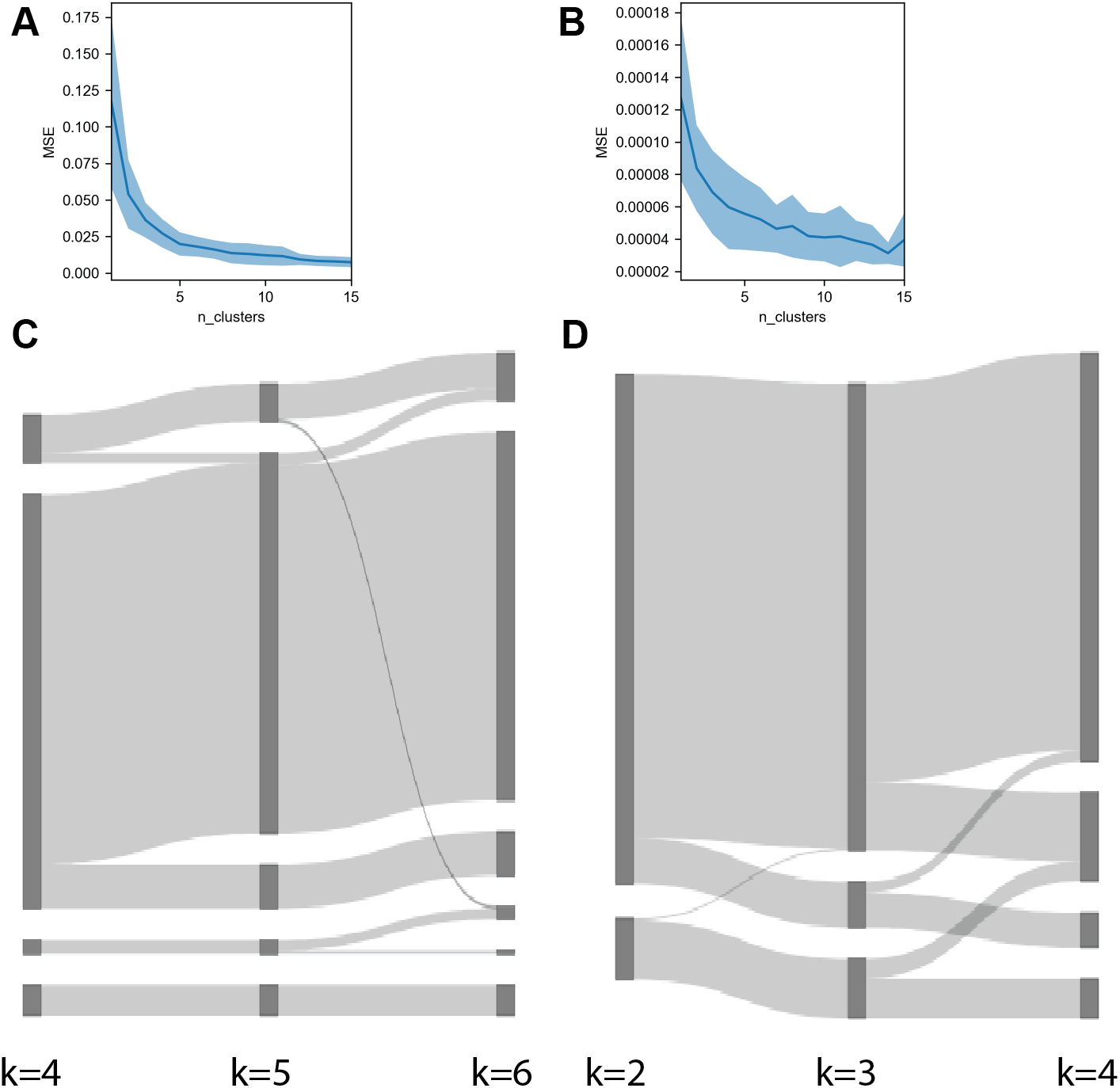
Selecting the number of cause categories for CFL (see Figures 5 and 6 for more details). (A) 10-fold cross-validated mean squared error in predicting E[*P* (*Y |X*)] from cause category assignment for different numbers of categories *k*, for the language-visuospatial (Cohort 2) analysis. (B) Same as A but for the BDI (Cohort 3) analysis. (C) Category assignments across values of *k* for the language-visuospatial (Cohort 2) analysis. (D) Same as C but for the BDI (Cohort 3) analysis.

**Figure S7:**
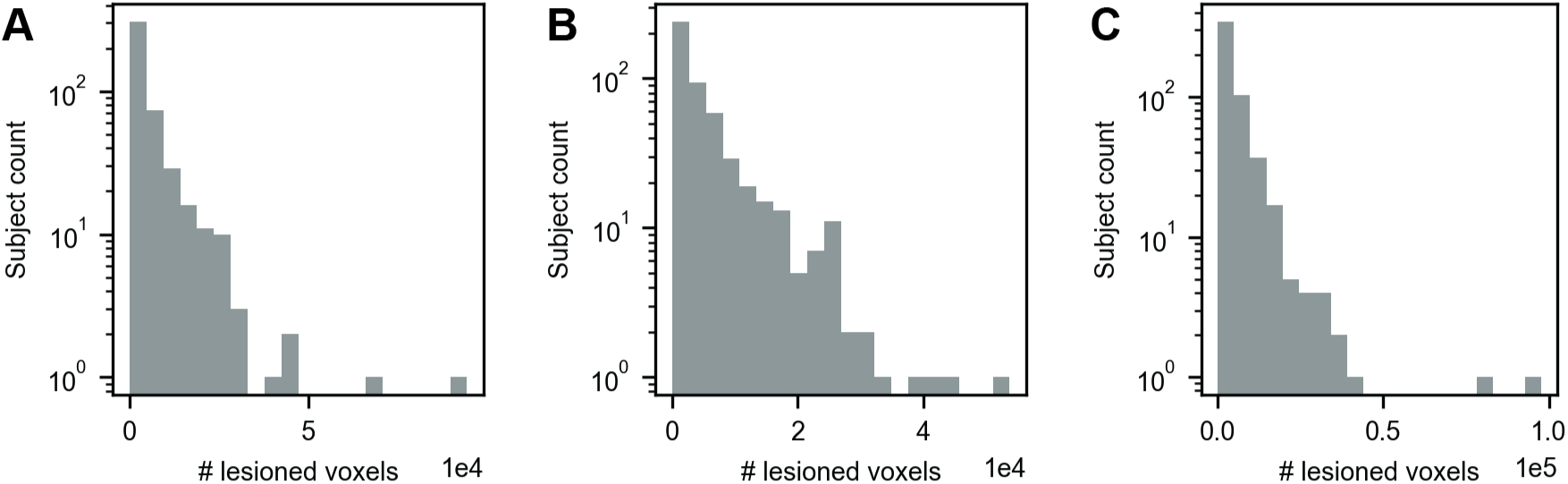
Lesion size distributions for (A) simulated cohort 1, (B) cohort 2, and (C) cohort 3.

**Figure S8:**
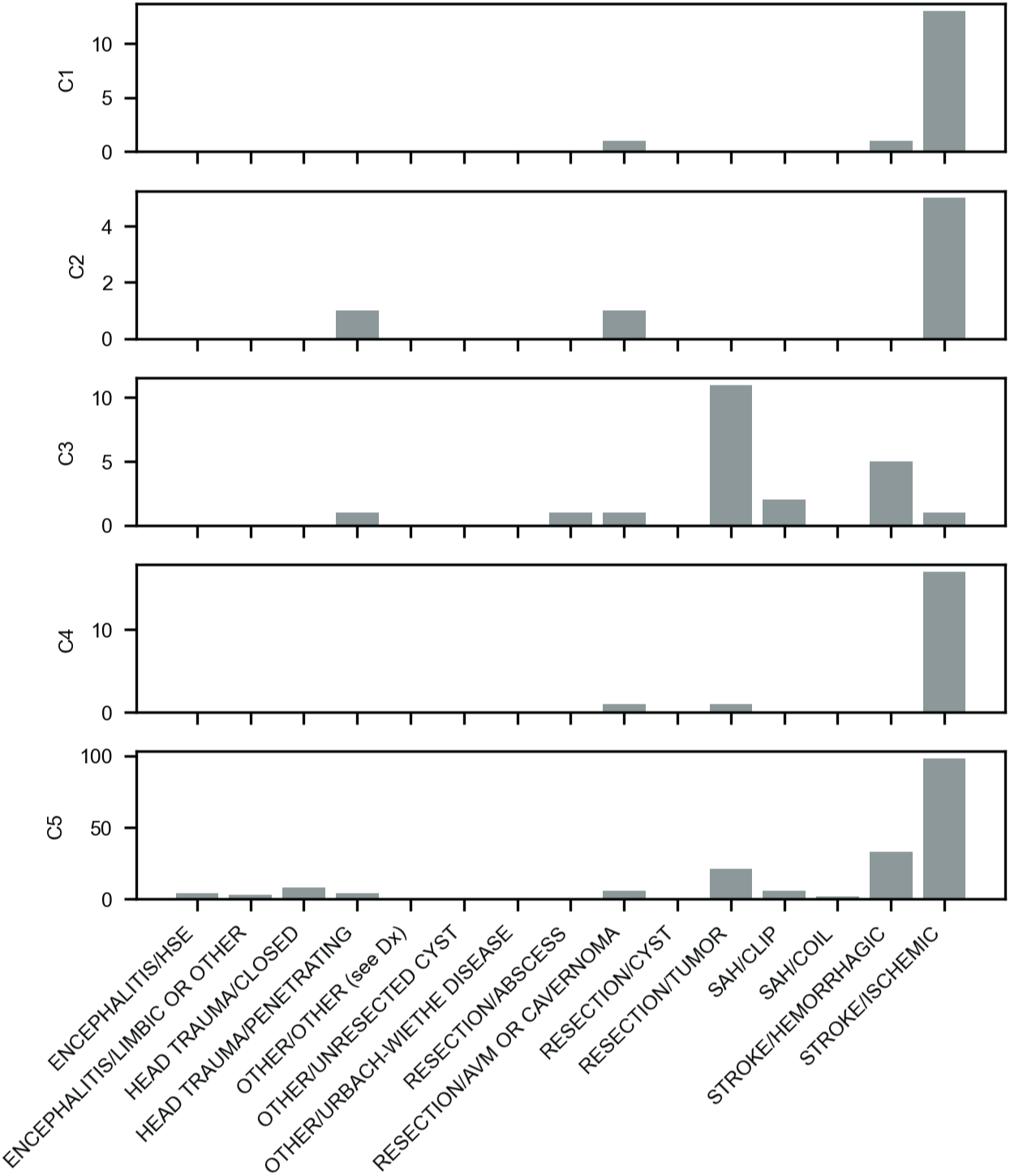
Lesion etiology distributions per CFL lesion category in language-visuospatial analysis. See Figure 5 for more details.

**Figure S9:**
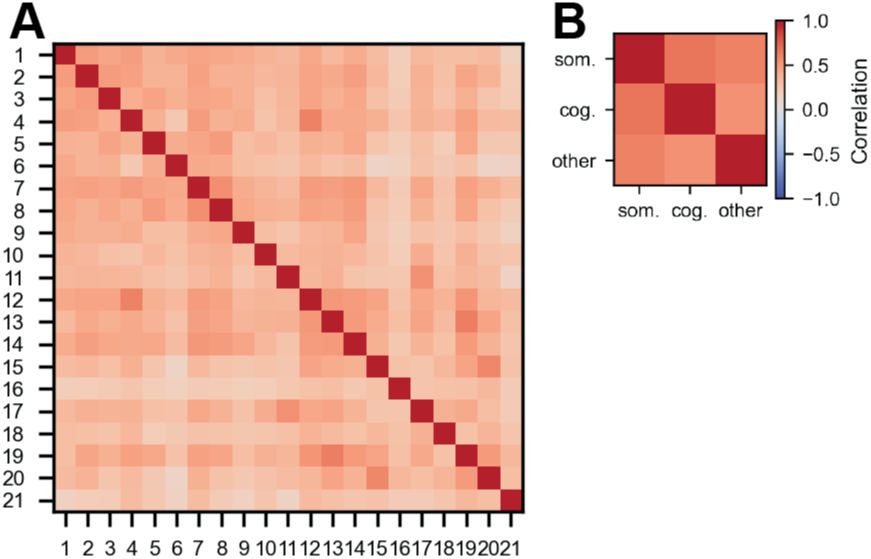
A) Pearson correlation for the 21 BDI items in Cohort 3. B) Pearson correlation matrix of the somatic and cognitive-affective BDI sub-scores. The list of BDI question labels and question assignments to somatic and cognitive-affective categories can be found in Figure 7.

**Figure S10:**
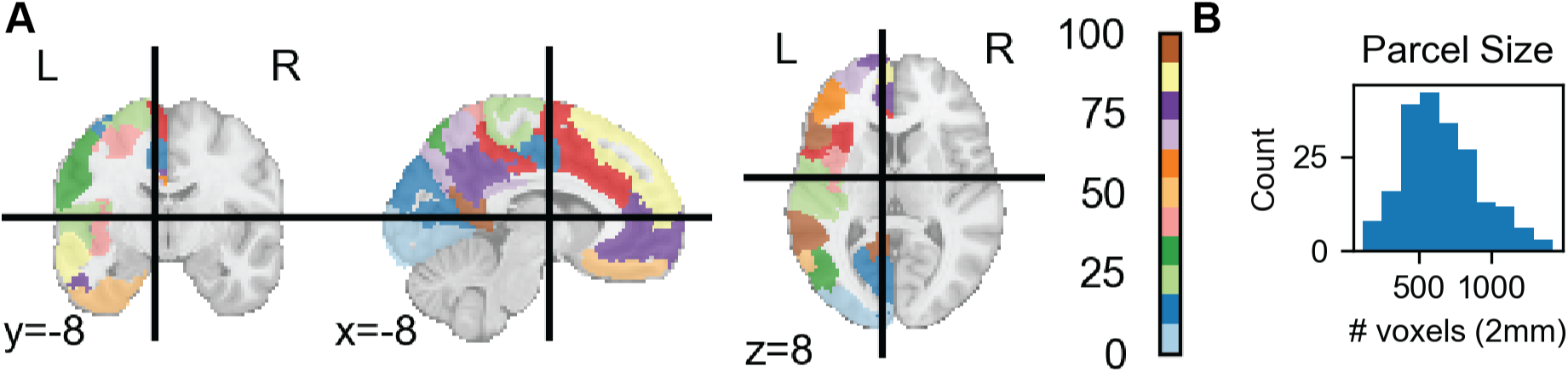
Simulated dataset parcellation (Cohort 1, see Figure 3 for more details). A) Left-hemispheric parcels in Schaefer-200 parcellation. B) Parcel size distribution.

**Figure S11:**
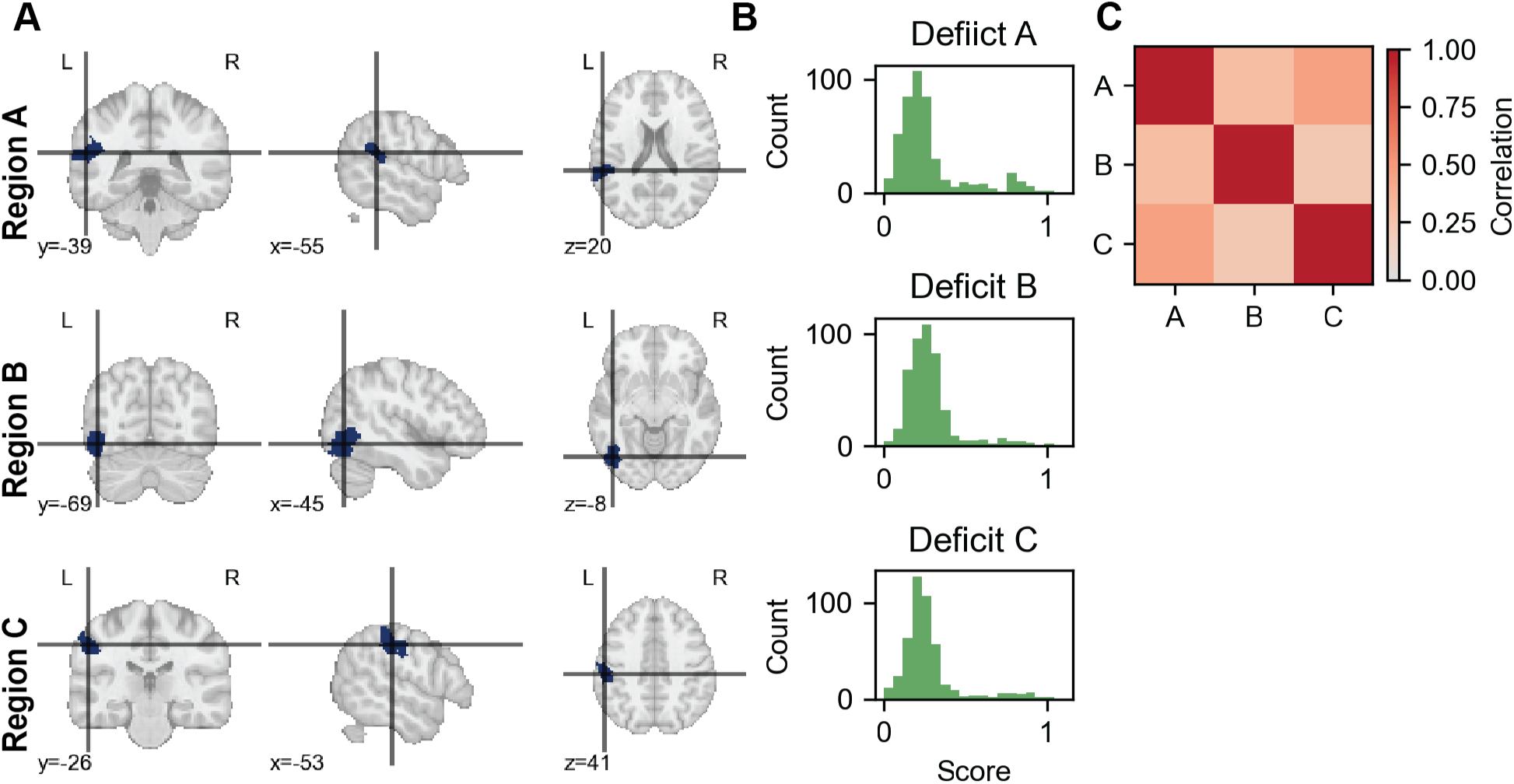
Simulated dataset (Cohort 1, see Figure 4 for more details). A) Three ground truth deficit-generating regions from the Schaefer-200 parcellation. B) Simulated deficit distributions. Deficits generated as a function of lesion overlap with the regions in A. C) Deficit correlations.

**Figure S12:**
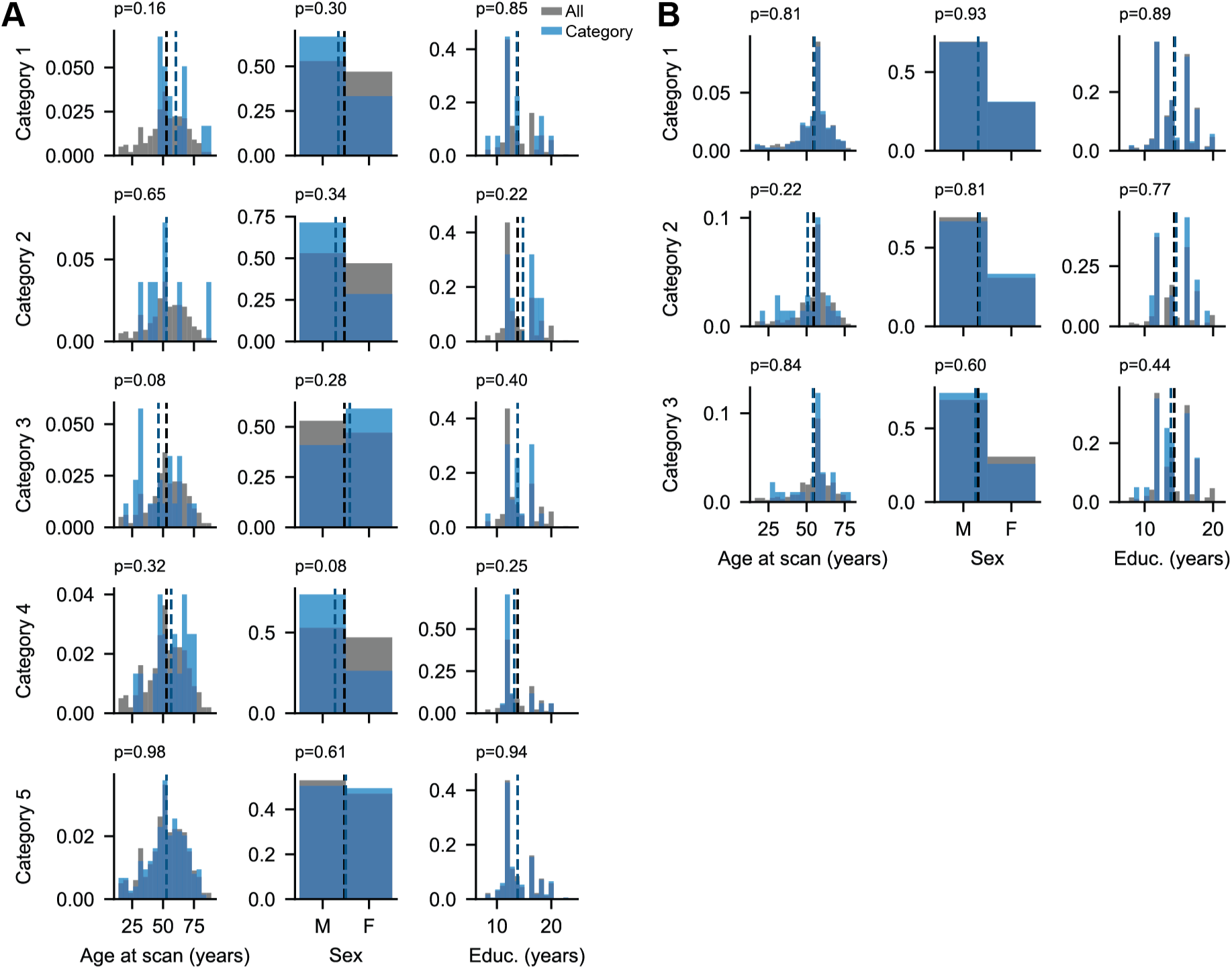
Demographic distributions by CFL lesion category for A) Cohort 2 and B) Cohort 3 analyses. Available demographics included: age at time of lesion scan, biological sex, and years of education. Full cohort distributions are shown in gray. Category-specific distributions are shown in blue. Distribution means are shown in dark gray (full cohort) and blue (category-specific). Normalized histogram densities, as opposed to raw counts, are shown, as most CFL categories are much smaller than the full population. P-values from Two-sided Mann-Whitney U test comparing category-specific to full cohort distributions are noted in the top left of each panel. All p-values are > 0.05 before multiple comparisons corrections.

**Figure S13:**
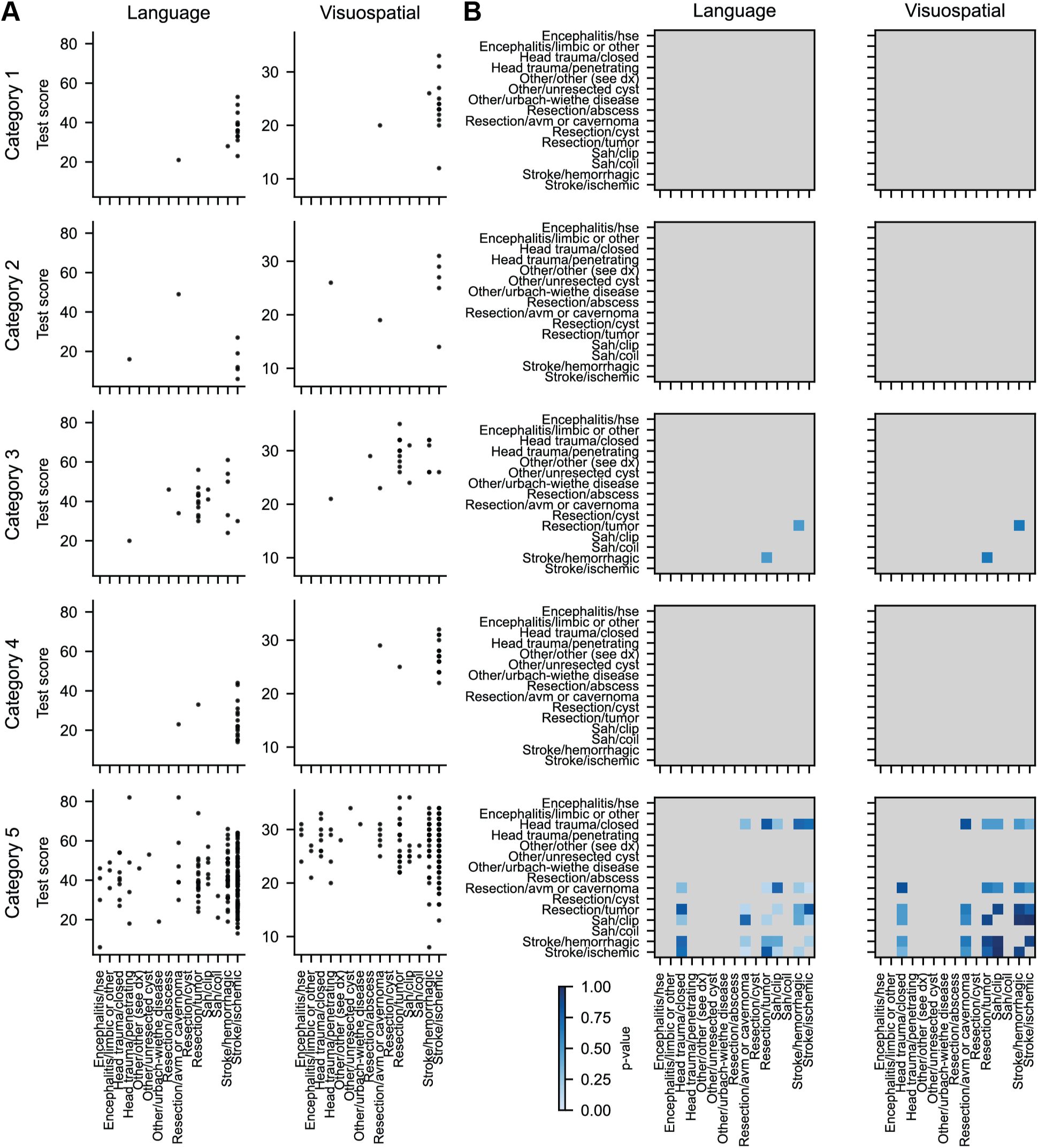
Relationship between lesion etiology and test scores per CFL lesion category for Cohort 2. A) Language (left) and visuospatial (right) scores per CFL lesion category (rows) and lesion etiology category (x-axis). B) Two-sided t-test p-values between pairs of lesion etiology groups for each CFL lesion category (rows) and test score (columns). Only lesion etiology categories with more than five subjects per lesion category and test score were compared (blue color map). Comparisons with less than five subjects are shown in gray. All p-values are > 0.05 before multiple comparisons corrections.

## Notes

### Competing Interest Statement

The authors declare no competing interests. Of note, A.D.B. is an unpaid co-founder of NeuroPred, Inc., a start-up company focused on lesion-informed stroke prognostication.

### Summary of Updates

Corrected typos in main text; Categories in Figure 7 re-ordered for consistency with Figure 6.

